# Genome-wide analysis of transcriptional bursting-induced noise in mammalian cells

**DOI:** 10.1101/736207

**Authors:** Hiroshi Ochiai, Tetsutaro Hayashi, Mana Umeda, Mika Yoshimura, Akihito Harada, Yukiko Shimizu, Kenta Nakano, Noriko Saitoh, Hiroshi Kimura, Zhe Liu, Takashi Yamamoto, Tadashi Okamura, Yasuyuki Ohkawa, Itoshi Nikaido

## Abstract

Transcriptional bursting is stochastic activation and inactivation of promoters, leading to discontinuous production of mRNA, and is considered to be a contributing factor to cell-to-cell heterogeneity in gene expression. However, it remains elusive how the kinetic properties of transcriptional bursting (*e.g.*, burst size, burst frequency, and noise induced by transcriptional bursting) are regulated in mammalian cells. In this study, we performed a genome-wide analysis of transcriptional bursting in mouse embryonic stem cells (mESCs) using single-cell RNA-sequencing. We found that the kinetics of transcriptional bursting was determined by a combination of promoter and gene body binding proteins, including polycomb repressive complex 2 and transcription elongation-related factors. Furthermore, large-scale CRISPR-Cas9-based screening and functional analysis revealed that the Akt/MAPK signaling pathway regulated bursting kinetics by modulating transcription elongation efficiency. These results uncover key molecular mechanisms underlying transcriptional bursting and cell-to-cell gene expression noise in mammalian cells.

## Introduction

Single gene expression in unicellular organisms displays significant cell-to-cell variations (*1*); a phenomenon called gene expression noise. It is thought that gene expression noise generates significant phenotypic diversity in unicellular organisms, improving the fitness of the species by hedging against sudden environmental changes (*2*). Such heterogeneity in gene expression is also observed in multicellular organisms during viral response and differentiation (*3*). It is widely believed that there are 2 orthogonal sources of gene expression noise: 1) intrinsic noise associated with stochasticity in biochemical reactions (*e*.*g*., transcription, translation); and 2) extrinsic noise induced by true cell-to-cell variations (*e*.*g*., differences in the microenvironment, cell size, cell cycle phase, and concentrations of cellular components) (*4, 5*).

It was well-established that the first step in gene expression, transcription, occurs with widely different bursting kinetics, as the promoter stochastically switches between “ON” and “OFF” states (Fig. 1A), while mRNAs are produced only during the ON state. This phenomenon is called transcriptional bursting and has been observed in various gene systems in a wide range of organisms (*5-8*). Since switching of the promoter state is stochastic in nature, transcriptional bursting induces heterogeneity in expression even between the two gene alleles in the diploid genome (Fig. 1B, C). This type of noise is considered to be a major part of intrinsic noise at the transcript level (*7, 9, 10*). Here, we refer to it as transcriptional bursting-induced noise (TBi noise, 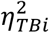). The mean number of mRNAs (*μ*) and TBi noise can be determined by the frequency of the promoter being in the active state (burst frequency, *f*), along with the number of transcripts produced per burst (burst size, *b*), and their degradation rate (*γ_m_*) (Fig. 1A, see Material and Methods) (*9, 11*).

**Fig. 1.**
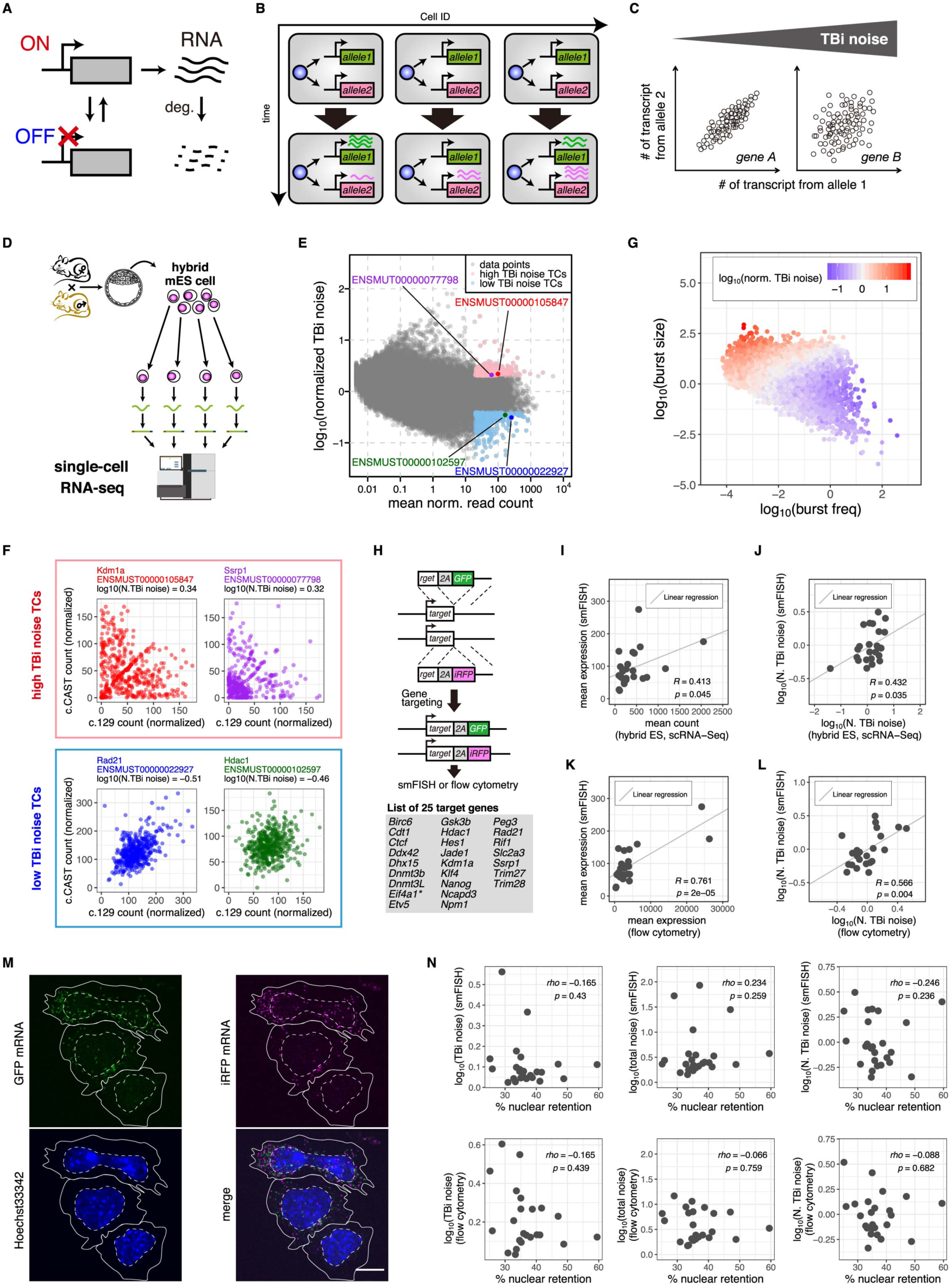
Genome-wide analysis of transcriptional bursting kinetics. **(A)** Upper panel shows a schematic representation of gene expression accompanied by stochastic switching between transcription permissive active (ON) and inactive (OFF) states (transcriptional bursting). **(B, C)** Transcriptional bursting induces not only inter-allelic but also cell-to-cell heterogeneity in gene expression. **(B)** If there are several cells with the same cellular state, they will show heterogeneity in gene expression among cells and even among alleles due to transcriptional bursting. **(C)** Scatter plots of the number of transcripts derived from individual alleles. Each data point indicates the expression data from a single cell. The larger the TBi noise, the more likely the distribution is to extend perpendicular to the diagonal. **(D)** Schematic representation of scRNA-seq using hybrid mouse embryonic stem cells (mESCs). **(E)** Scatter plot of mean normalized read counts and normalized TBi noise of individual transcripts (TCs) revealed by scRNA-seq. **(F)** Representative scatter plots of normalized individual allelic read counts of high and low TBi noise TCs. Colors of spots are corresponding to **(E)**. N. TBi noise: normalized TBi noise. **(G)** Scatter plot of burst size and burst frequency of individual TCs. Color code represents the amplitude of the normalized TBi noise. **(H)** Schematic representation of knock in of GFP and iRFP gene cassette into individual alleles of mESC derived from inbred mice. Lower panel represents a list of targeted genes. Only for a gene with an asterisk, knock-in (KI) cassettes were inserted immediately downstream of the start codon. For the other target genes, KI cassettes were inserted immediately upstream of the stop codon. **(I)** Scatter plot of mean number of TCs of targeted genes in KI cell lines counted by smFISH and mean normalized read counts of corresponding genes in hybrid mESCs revealed by scRNA-seq. **(J)** Scatter plot of normalized TBi noise of TCs of targeted genes in KI cell lines revealed by smFISH and that of corresponding genes in hybrid mESCs revealed by scRNA-seq. **(K)** Scatter plot of mean expression levels of targeted genes in KI cell lines revealed by flow cytometry and mean number of TCs of targeted genes in KI cell lines counted by smFISH. **(L)** Scatter plot of normalized TBi noise of targeted genes in KI cell lines revealed by flow cytometry and that of targeted genes in KI cell lines counted by smFISH. **(M)** Representative images of smFISH using GFP and iRFP probes in *Nanog* KI cell line. Solid and dashed lines represent the plasma and nuclear membranes, respectively. Scale bar: 10 µm. **(N)** Scatter plots of nuclear retention rate of mRNA and either TBi noise, total noise or normalized TBi noise revealed by smFISH or flow cytometry.

Interestingly, although it has been proposed that the suppression of promoter reactivation is essential for the control of transcriptional bursting, the underlying mechanism remains largely unknown (*12*). However, it has been demonstrated in yeast that *cis*-regulatory elements (such as the TATA box) and chromatin accessibility at the core promoter regulate transcriptional bursting kinetics (*10*). In higher eukaryotes, kinetic properties of transcriptional bursting have also been extensively studied by using smFISH, MS2 system, and destabilized reporter proteins (*7, 13-15*). These reports indicate that mammalian genes are transcribed with widely different bursting kinetics (burst sizes and frequencies). Interestingly, it was shown that transcriptional bursting can be influenced by local chromatin environments as the same reporter-gene displays distinct bursting kinetics when inserted into distinct genomic locations (*11*). Despite these studies, we still lack a comprehensive understanding of how transcriptional bursting is regulated at the molecular level.

Mouse embryonic stem cells (mESCs) are derived from the inner cell mass of preimplantation embryo. A large number of genes in mESCs, cultured on gelatin in standard (Std) medium containing serum and leukemia inhibitory factor (LIF), showed cell-to-cell heterogeneity in expression (*16*). For example, several genes encoding key transcription factors, including *Nanog*, displayed heterogeneous expression levels in the inner cell mass and mESCs (*17*). It was proposed that the heterogeneity in gene expression breaks the symmetry within the system and primes cells for subsequent lineage segregation (*16*). Previously, we quantified *Nanog* transcriptional bursting kinetics in live cells using the MS2 system and found that TBi noise is a major cause of heterogeneous NANOG expression in mESCs (*18*). A recent study using intron-specific smFISH revealed that most genes in mESCs are transcribed with bursting kinetics (*19*).

In the present study, to investigate the extent to which TBi noise contributes to heterogeneous gene expression in mESCs, we performed single-cell RNA-seq (RamDA-seq) (*20*) using hybrid mESCs for a comprehensive, genome-wide analysis of the TBi noise, burst size, and frequency. Genomic analysis and functional validation revealed that transcriptional bursting kinetics were related to a combination of promoter and gene body binding proteins, including polycomb repressive complex 2 and transcription elongation-related factors. In addition, CRISPR library screening revealed that the Akt/MAPK pathway regulates transcriptional bursting via modulating the transcription elongation efficiency. These results reveal new molecular mechanisms underlying transcriptional bursting and TBi noise in mammalian cells.

## Results

### Measuring transcriptional bursting-induced noise in hybrid mESCs with single-cell RNA-seq

To study the genome-wide kinetic properties of transcriptional bursting, we analyzed allele-specific mRNA levels from 447 individual 129/CAST hybrid mESCs (grown on Laminin-511 [LN511] without feeder cells in the G1 phase) by single-cell (sc) RamDA-seq—a highly sensitive RNA-seq method (*20*) (Fig. 1D, S1A, S2A-E). Some genes have transcript variants with different transcription start sites (TSSs). Since the kinetic properties of transcriptional bursting may differ depending on the promoter, we mainly used transcript (TC)-level abundance data rather than gene-level abundance data to estimate the kinetic properties of transcriptional bursting (see Material and Methods, Fig. S1B-G). TBi noise, which is a major part of intrinsic noise at TC level (*7, 9, 10*), can be estimated from the distribution of the number of mRNAs produced by the two alleles (*4*) (see Material and Methods). Further, we normalized TBi noise based on the expression level and transcript length of the gene (Fig. 1E, S1B-G). We excluded low abundance transcripts (with read count of less than 20) from the downstream analysis, as it is difficult to distinguish whether technical or biological noise contributes to the measured heterogeneity of allele-specific expression. We ranked the genes based on their normalized TBi noise and defined the top and bottom 5% TCs as high and low TBi noise TCs, respectively (Fig. 1E). As expected, high TBi noise TCs showed larger inter-allelic expression heterogeneity than low TBi noise TCs (Fig. 1F, Table S1, S2).

Because the mRNA degradation rate can affect TBi noise (see Material and Methods), we next checked the relationship between the published mRNA degradation rate in mESCs (*21*) with the normalized TBi noise that we measured, but no correlation was observed (Fig. S1G). We next estimated the burst size and frequency of each TC based on the published mRNA degradation rate and the TBi noise (Fig. 1A, Fig. S2F-K, see Material and Methods) (*9, 11*). As expected, we found that TCs with larger burst sizes and lower burst frequencies tend to show higher normalized TBi noise and vice versa (Fig. 1G). Thus, the normalized TBi noise is positively well correlated with the ratio of the burst size to the burst frequency (Spearman rho = 0.869).

To confirm whether the TBi noise measured by scRNA-seq indicates true gene expression noise, we first selected 25 genes showing medium expression levels and diverse TBi noise levels. Then, using CRISPR-Cas9 genome editing, we integrated a GFP and an iRFP reporter gene separately into both alleles of these genes in an inbred mESC line (KI mESC lines) (Fig. 1H, S1H, S3). It is worth noting that the GFP and iRFP reporter cassettes were flanked by a 2A peptide and a degradation promoting sequence and was inserted immediately upstream of the stop codon of each allele (only one cell line, reporter cassettes were knocked in immediately downstream of the start codon, see Fig. 1H and S3). The 2A peptide separates the reporter protein from the endogenous gene product. The degradation promoting sequence ensured rapid degradation of GFP or iRFP reporter so that the amount of fluorescent protein produced in the cell would reflect the cellular mRNA levels. Using these cell lines, we measured the mean expression levels and normalized TBi noise for the 25 genes with smFISH and found that the two parameters showed a significant correlation with scRNA-seq based measurements (Fig. 1I, J, S1H, Table S3). Furthermore, flow cytometry analysis confirmed that the mean expression level and normalized TBi noise at the protein level also showed a significant correlation with the smFISH-based measurements (Fig. 1K, L, S1I). There was a substantial correlation between the expression level of the endogenous target protein and that of the knocked-in fluorescent protein in all tested genes (Fig. S1J, K). These validation experiments demonstrated the robustness of scRNA-seq in determining TBi noise and revealed that heterogeneity in the expression of the tested genes originated from the variation of the mRNA levels.

Recently, it has been reported that TBi noise can be buffered by nuclear retention of mRNA molecules (*22*). Here, we investigated the relationship between the subcellular localization of mRNA and normalized TBi noise in KI cell lines by smFISH (Fig. 1M). We observed no correlation between the mRNA nuclear retention rate and the TBi noise, total noise or the normalized TBi noise at the mRNA or protein levels (Fig. 1N). Thus, it is unlikely that mRNA nuclear retention plays a role in buffering TBi noise for the 25 genes tested in mESCs.

### Identifying molecular determinants of transcriptional bursting

It has been reported that promoters with the TATA box tend to show higher burst size and gene expression noise than promoters without it in yeast and mouse ES cells (*8, 10, 12, 23*). To confirm whether our data were consistent with those findings, we compared the transcriptional bursting kinetics of genes either with or without a TATA box. Although no significant difference was observed in the burst frequency, the burst size and normalized TBi noise were significantly higher in the promoters with the TATA box than promoters without it (Fig. 2A-C). These data, which are consistent with previous reports, validate the quality of our results and support the involvement of the TATA box in burst size and gene expression noise (Fig. 2A-C).

**Fig. 2.**
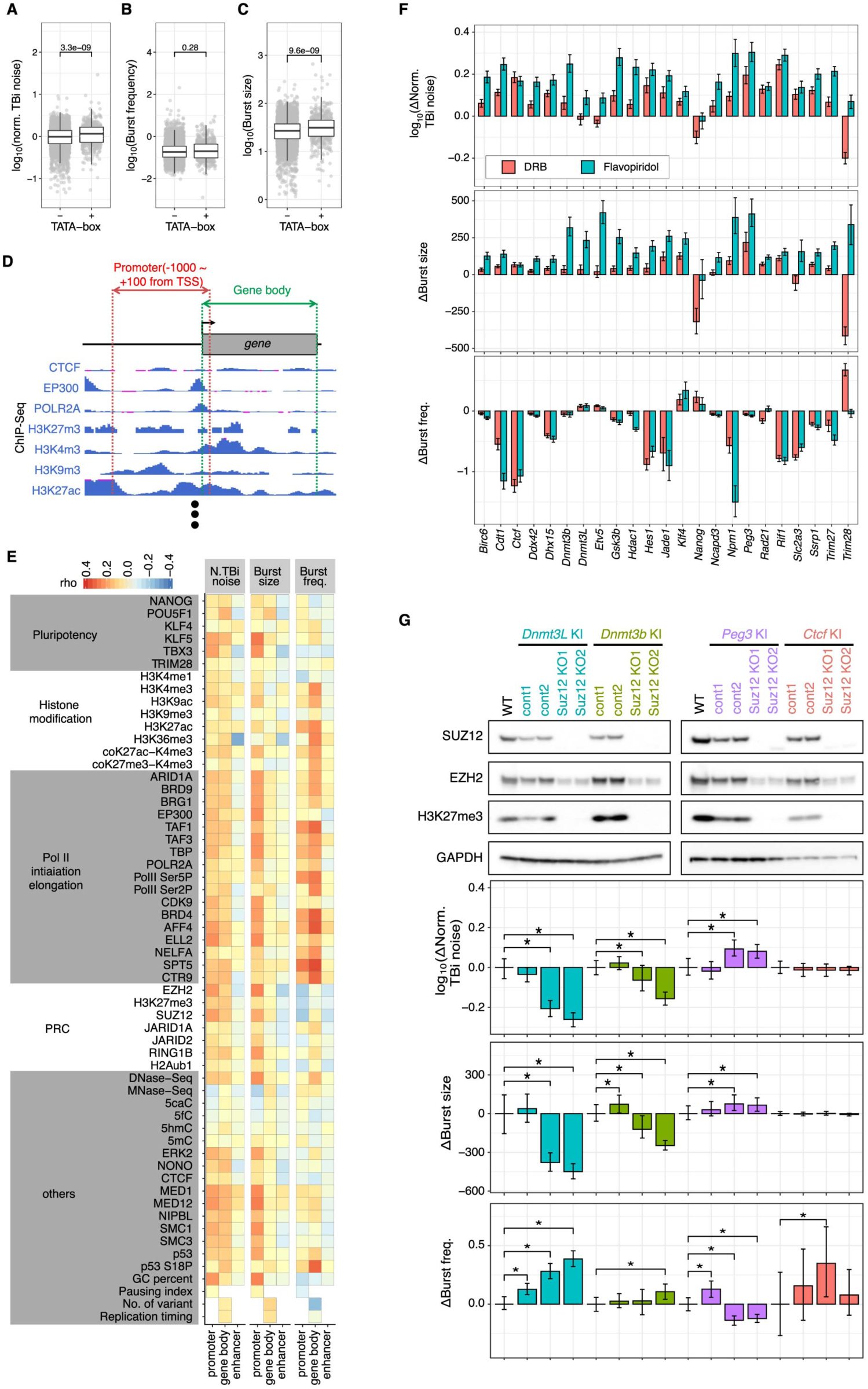
PRC2 and transcriptional elongation are involved in transcriptional bursting kinetics. **(A-C)** Normalized TBi noise and burst size of genes with TATA box are significantly higher than those of genes without TATA box. **(D)** Schematic representation of calculating reads per million (RPM) at the promoter and gene body from ChIP-seq data. In addition, similar calculations were also performed for enhancers (see Methods). **(E)** Heatmaps of Spearman’s rank correlation between promoter, gene body or enhancer-associated factors and either normalized TBi noise, burst size or burst frequency. **(F)** Effect of transcription elongation inhibitor DRB and Flavopiridol treatment on transcriptional bursting kinetics. The KI cell lines were treated with DRB or Flavopiridol for 2 days, analyzed by flow cytometry, and transcriptional bursting kinetics were calculated. Δnormalized TBi noise, Δburst size and Δburst frequency are residuals of normalized TBi noise, burst size and frequency of inhibitor-treated cells from that of control cells, respectively. Error bars indicate 95% confidence interval. **(G)** Effect of *Suz12* K/O on normalized TBi noise. *Suz12* K/O cell lines derived from *Dnmt3l*, *Dnmt3b*, *Peg3*, and *Ctcf* KI cell lines were established. Upper panel represents the result of western blotting. Compared with the control cell lines (cont1 and 2), the *Suz12* KO cell lines (Suz12 KO1 and 2) showed complete loss of the SUZ12 and H3K27me3 signals and a decrease in EZH2 levels. In the lower part of the panel, the Δnormalized TBi noise, Δburst size and Δburst frequency compared with the control (cont1) are shown. Error bars indicate 95% confidence interval. Asterisks indicate significance at *P* < 0.05.

Next, we compared the kinetic properties of transcriptional bursting to genome-wide transcription factor binding patterns (Fig. 2D, see Material and Methods). Specifically, we calculated Spearman’s rank correlations between the kinetic properties of transcriptional bursting and ChIP-seq enrichment in the promoter, gene body, or enhancer elements (Fig. 2E). We found that the localization of several transcription regulators (*e.g.*, EP300, ELL2, and MED12) in the promoter showed substantial positive correlations with the burst size. However, the correlation coefficients between the burst size and transcription regulators bound in enhancers were overall relatively low. This is consistent with the reports that burst size is mainly controlled by the promoter region (*8*).

On the other hand, it has been suggested that the distal enhancers are important for regulating burst frequency (*24*). We found that the localization of several factors (*e.g.*, BRD9, BRD4, and CTR9) in enhancers showed relatively lower, but positive correlations with the burst frequency.

Interestingly, we found that the localization of transcription elongation factors (*e.g.*, H3K36me3, BRD4, AFF4, SPT5, and CTR9) on the gene body was positively correlated with the burst frequency (Fig. 2E). It has been shown that, after transcription initiation, the RNA Pol II could pause near the promoter region before transitioning to the productive elongation stage (*25*); we found that the degree of RNA pol II pausing measured by the pausing index has a negative correlation with the burst frequency (Fig. 2E), consistent with a model that RNA pol II pausing might suppress the bursting frequency.

To dissect the link between transcription elongation and burst frequency, we inhibited positive transcription elongation factor (P-TEFb) with DRB and Flavopiridol in KI mESC lines cultured in 2i conditions (*26*). 2 days after DRB and Flavopiridol treatment, cells were subjected to flow cytometry analysis (Fig. S4A, B). We found that DRB and Flavopiridol treatment increased the normalized TBi noise and burst size in most of the cell lines (Fig. 2F). However, the effects of DRB and Flavopiridol treatment on the burst frequency were highly gene-specific, suggesting that transcriptional elongation likely contributes to the regulation of both burst size and frequency, while the regulation of the burst frequency by transcriptional elongation is more context dependent.

The promoter localization of PRC2 subunits (EZH2, SUZ12, and JARID2) was negatively correlated with the burst frequency but positively correlated with the burst size and normalized TBi noise (Fig. 2E), suggesting a possible link between PRC2 and TBi noise. To test how PRC2 regulates transcriptional bursting, we inactivated PRC2 functionality by knocking out of SUZ12 (*27*) in *Dnmt3l*, *Dnmt3b*, *Peg3*, and *Ctcf* KI cell line (Fig. 2G). These targeted genes showed relatively high H3K27me3 enrichment at the promoter compared to the other available KI-targeted genes. Loss of H3K27me3 modification in *Suz12* knockout (K/O) cell lines was confirmed by western blotting (Fig. 2G). Then, we quantified GFP and iRFP expression levels by flow cytometry in the *Suz12* K/O and control cell lines. We found that normalized TBi noise and burst size of *Dnmt3l* and *Dnmt3b* were significantly reduced by *Suz12* K/O (Fig. 2G). In contrast, *Suz12* K/O significantly increased normalized TBi noise and burst size of *Peg3*; but no significant change was observed for *Ctcf*. The burst frequency of *Dnmt3l* was significantly increased while that of *Peg3* was dramatically reduced by *Suz12* K/O. These results suggest that the control of the transcriptional bursting kinetics by PRC2 may also be context-dependent.

### Transcriptional bursting is regulated by a combination of promoter and gene body associated factors

To study the combinatorial regulations underlying transcriptional bursting kinetics, we first classified genetic and epigenetic features based on the sequence and TF binding patterns at the promoter and gene body of high TBi noise TCs into 10 clusters (Fig. 3). Then, to identify the features that most contributed to distinguishing a cluster of high-TBi noise TCs from low-TBi noise TCs, we performed OPLS-DA modeling, which is a useful method for identifying features that contribute to class differences (*28*). In 8 out of the 10 clusters, the model successfully separated the high TBi noise TCs from the low TBi noise TCs (Fig. 3). Specifically, we obtained the top three positively and negatively contributing factors by using S-plot (Fig. 3). For example, in cluster 3, the promoter binding of PRC2-related factors were positive contributors to the TBi noise while the gene body localization of TAF1, BRD4, CTR9 were negative contributors. This result suggests that the promoter localization of PRC2-related factors influences bursting properties in a gene specific manner.

**Fig. 3.**
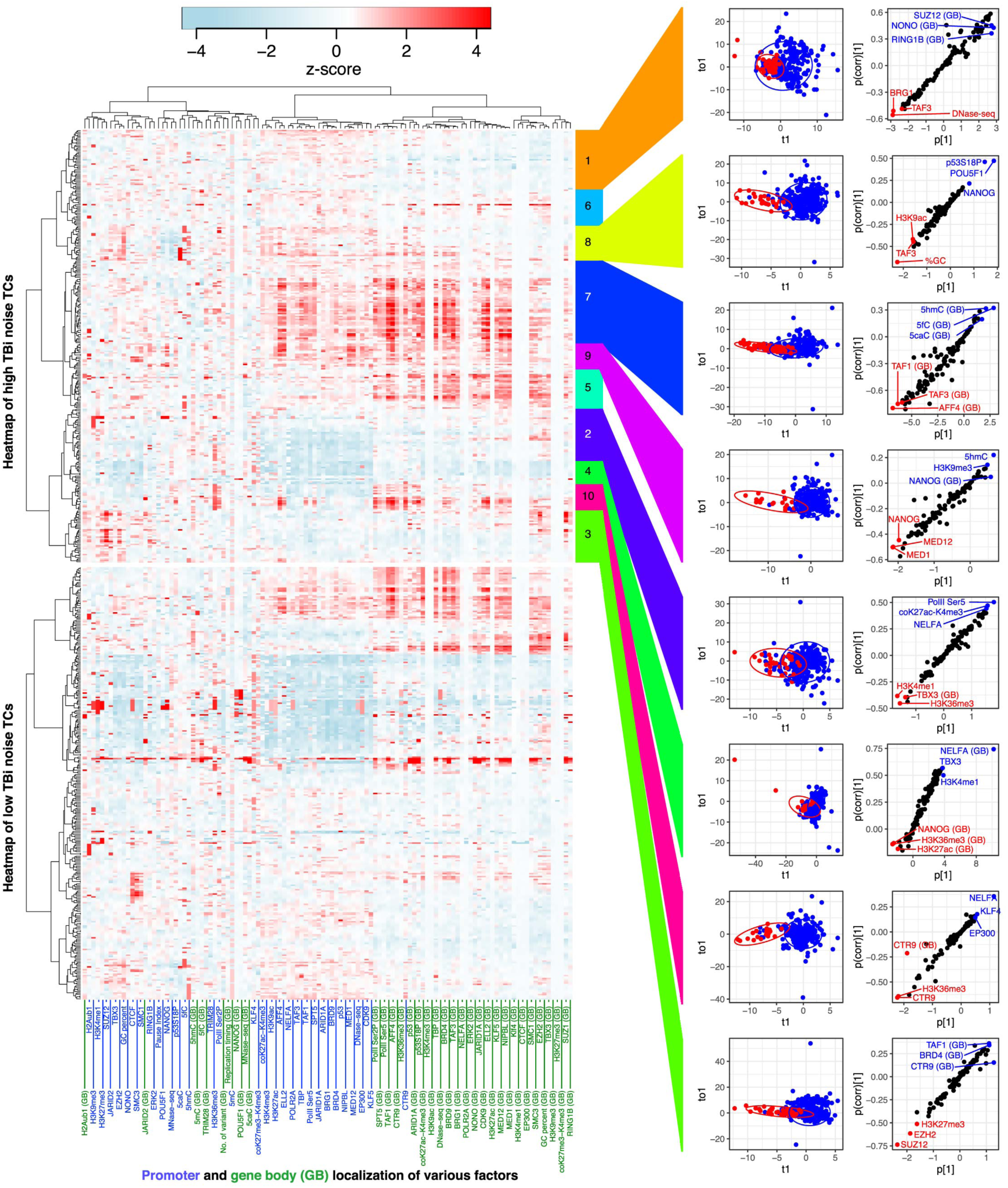
Normalized TBi noise is determined by combinations of promoter and gene body associating factors. The left side of the panel shows a heat map of promoter and the gene body (GB) localization of various factors with high and low TBi noise transcripts (TCs). The high TBi noise TCs were classified into 10 clusters, and each cluster of high TBi noise TCs and low TBi noise TCs were subjected to OPLS-DA modeling. The right side of the panel represents score plots of OPLS-DA and S-plots constructed by presenting covariance (*p*) against correlation [*p*(*corr*)].

In cluster 10, promoter localization of H3K36me3 - a histone mark associated with transcriptional elongation, and promoter and gene body localization of CTR9 - a subunit of PAF1 complex involved in transcriptional elongation were positive contributors, while promoter localization of NELFA was a strong negative contributor (Fig. 3). These results imply that transcriptional elongation is involved in the regulation of normalized TBi noise in this cluster. In addition, we used a similar analysis to identify factors regulating burst size and frequency and found that these two kinetic properties were also determined by a combination of promoter and gene body-associated factors (Fig. S5). Collectively, these results suggest that the kinetic properties of transcriptional bursting in mammalian cells were regulated by a combinatory suite of promoter and gene body binding factors in a context-dependent manner.

### Genome-wide CRISPR library screening identified genes involved in regulation of TBi noise

To identify genes regulating TBi noise in an unbiased manner, we next performed high-throughput screening with the CRISPR knockout library (*29*). Specifically, the CRISPR lentiviral library targeting genes in the mouse genome was introduced into *Nanog*, *Dnmt3l*, *Trim28* KI cell lines. Although genes with high TBi noise showed a larger variation in the expression levels of one allele (*e*.*g*., GFP) and the other allele (*e*.*g*., iRFP) perpendicular to the diagonal line (Fig. 1C, F), we found that the loss of genomic integrity (*e*.*g*., by a loss of function of p53) induced instability in the number of alleles, resulting in an unintended increase in TBi noise levels in a pilot study. Therefore, to reduce false negatives and to selectively enrich cell populations with suppressed TBi noise, we first sorted out cells showing expression levels close to the diagonal line of GFP and iRFP expression by FACS (Fig. 4A). After expanding the sorted cells for 1 week, the cells were sorted again. Such sorting and expansion procedures were repeated 4 times in total to selectively enrich cell populations with suppressed TBi noise. Even in the genes showing high TBi noise, a large fraction of cells showed a smaller variation in the expression levels of one (*e*.*g*., GFP) and other alleles (*e*.*g*., iRFP) perpendicular to the diagonal line (Fig. 1C, F). Therefore, the enrichment of the cells with low TBi noise by these repeated-sorting procedures could reduce false positives. Finally, we compared the targeted K/O gene profile in the sorted cells with that in unsorted control by high-throughput genomic DNA sequencing (Fig. 4A). To gain a comprehensive picture of genes involved in TBi noise regulation, we performed KEGG pathway analysis of the enriched targeted genes (top 100) and depleted targeted genes (bottom 100) in the three cell lines (Fig. 4B, C). Strikingly, we found that mTOR and MAPK signaling pathways were involved in promoting TBi noise in all three cell lines (Fig. 4C). Previous studies demonstrated that “Proteoglycans in Cancer” and “Sphingolipid signaling” pathways include mTOR and MAPK pathways could crosstalk with each other via the PI3/Akt pathway (Fig.4D) (*30*). To test whether these signaling pathways are involved TBi noise regulation, we conditioned *Nanog*, *Trim28*, *Dnmt3l* KI cells with inhibitors for the MAPK, mTOR, and Akt pathways (Fig. 4D-F). We found that when treated with the Akt inhibitor MK-2206 alone, normalized TBi noise decreased in all three cell lines (Fig. 4F). Treatment with MK-2206 and MEK inhibitor PD0325901 (PD-MK condition) resulted in a substantial decrease in the normalized TBi noise in all cell lines. In addition, normalized TBi noise was reduced in most KI cell lines under the PD-MK condition (Fig. 4G) while mRNA degradation rates were largely unaltered (Fig S6). Furthermore, there were 1 and 3 genes showing log_10_ (Δnormalized TBi noise) more than −0.05 in cells cultured in the PD-MK and 2i conditions, respectively (Fig. 4G), suggesting that more genes showed reduced normalized TBi noise under the PD-MK condition than the 2i condition. It is worth noting that, upon PD-MK treatment, the burst size decreased while the burst frequency increased for most genes compared with the untreated condition. However, the degree of changes varied between different genes. This suggests that the decrease in normalized TBi noise under the PD-MK condition is likely caused by the changes in both burst size and burst frequency.

**Fig. 4.**
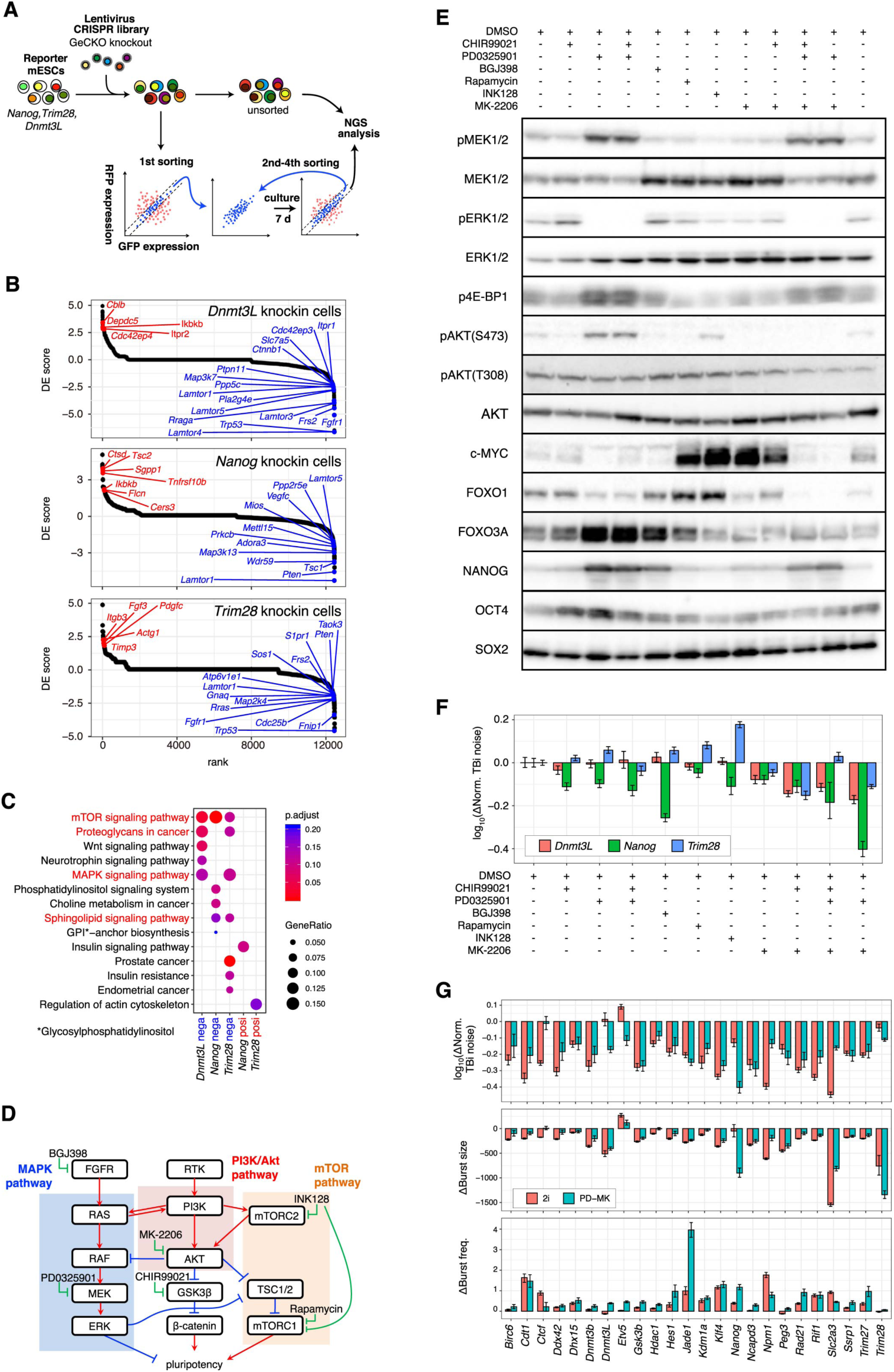
CRISPR library screening of genes involved in TBi noise regulation. (A) **Schematic** diagram of CRISPR lentivirus library screening. Screening was performed independently for each of the three (*Nanog*, *Trim28*, *Dnmt3l*) KI cell lines. **(B)** Ranked DE score plots obtained by performing CRISPR screening on three cell lines. The higher the DE score, the more the effect of enhancing TBi noise. While, the smaller the DE score, the more likely the gene can suppress the TBi noise. **(C)** KEGG pathway analysis. KEGG pathway analysis was performed with the upper or lower 100 genes of DE score obtained from the CRISPR screening (referred as posi and nega, respectively). The pathways shown in red indicate hits in multiple groups of genes. Genes corresponding to these pathways are labeled in **(B)**. See also Fig. S6 for these pathways. **(D)** A simplified diagram of MAPK, Akt, mTOR signaling pathways. These pathways are included in the pathways highlighted in red in **(C)**, and crosstalk with each other. **(E)** Western blot of cells treated with signal pathway inhibitors. **(F)** Δnormalized TBi noise of cells treated with signal pathway inhibitors against control (DMSO treated) cells. Error bars indicate 95% confidence interval. **(G)** Twenty-four KI cell lines were conditioned to 2i or PD-MK conditions and subjected to flow cytometry analysis. Δnormalized TBi noise, Δburst size and Δburst frequency against control (DMSO treated) cells are shown. Error bars indicate 95% confidence interval.

It was shown that Akt or MAPK pathways promote cell cycle progression (*30*). As expected, we found that PD-MK treatment significantly reduced the proliferation rates of mESCs (Fig. 5A). However, we did not observe increased cell apoptosis after the PD-MK treatment (Fig. 5B). The cell cycle distribution was also unaffected by the PD-MK treatment, (Fig. 5C), suggesting a uniformed slowdown of individual cell cycle phases in the PD-MK condition. However, the expression of pluripotent markers was largely unaffected under the PD-MK condition (Fig. 4E, 5D). More importantly, we can generate chimeric mice using mESCs cultured in the PD-MK conditions, suggesting that PD-MK treatment does not affect mESC pluripotency (Fig. 5E).

**Fig. 5.**
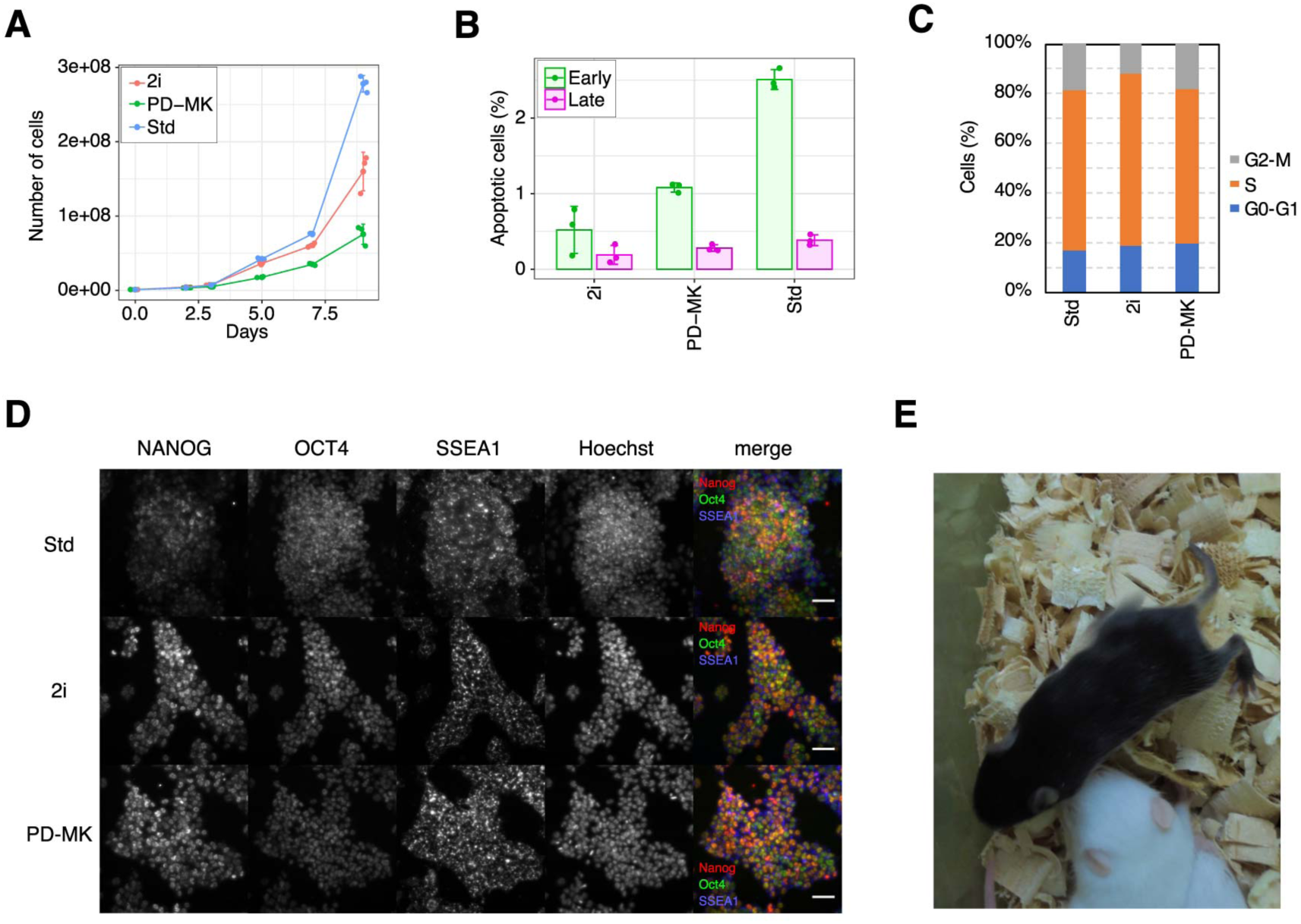
PD-MK conditioned mESCs retain pluripotency. **(A)** Growth curve of mESC conditioned to Std, 2i, PD-MK conditions. Error bars indicate standard deviation, *n* = 3. **(B)** Percentage of apoptotic cells of mESC conditioned to Std, 2i, PD-MK conditions. Error bars indicate standard deviation, *n* = 3. **(C)** Cell cycle distribution of mESCs conditioned to Std, 2i and PD-MK conditions. **(D)** Immunofluorescence of pluripotency markers (NANOG, OCT4, SSEA1) of mESCs conditioned to Std, 2i, PD-MK conditions. The images show maximum intensity projections of stacks. Scale bar: 50 μm. **(E)** Chimeric mice with black coat color generated from C57BL6NCr ES cells conditioned to PD-MK condition and then to Std condition before injection into albino host embryos.

To further characterize how PD-MK culturing condition affects mESC gene expression programs, we performed RNA-Seq analysis of mESCs cultured in standard (Std), 2i and PD-MK conditions. Flow cytometry analysis showed that more genes showed a decrease in normalized TBi noise under the PD-MK condition than under the 2i condition (Fig. 4G). To obtain a comprehensive view of the genes involved in TBi noise suppression under the PD-MK condition compared to the 2i condition, we performed GO analysis and found that the “transcription elongation factor complex” related genes were significantly enriched in the upregulated genes in the PD-MK condition compared with the 2i condition (Fig. 6B, C). Furthermore, the upregulated genes are also enriched in factors involved in transcriptional regulation (Fig. 6B), consistent with our observation that there is a positive correlation between gene body localization of transcription elongation factor and the burst frequency (Fig. 2C). Thus, it is possible that upregulation of transcription elongation factors could promote transcriptional bursting frequency and thus reduce the TBi noise in the PD-MK condition. We also found that the expression of *Aff1* and *Aff4*, that encode subunits of super elongation complex (SEC) (*25*) was significantly upregulated under the PD-MK condition (Fig. 6C). Next, we treated cells cultured in the PD-MK condition with P-TEFb inhibitor Flavopiridol or AFF1/4 inhibitor KL-2 (*31*) for 2 days, and compared the TBi noise in these cells with control cells cultured in the PD-MK medium (Fig. 6D, S4C, D). We found that the normalized TBi noise was significantly increased in the majority of genes, but decreases were also observed in a small fraction of genes (Fig. 6D). We also examined how transcriptional elongation inhibitors would affect the burst size and frequency, and found that, for most of the genes, Δburst sizes were small, while burst frequency was substantially reduced overall. This suggests that PD-MK treatment enhances transcription elongation efficiency but does not strongly affect the burst size at least for genes tested here. Furthermore, since P-TEFb and SEC inhibitors affect Δburst frequency in a similar fashion for most genes analyzed here, it is likely that SEC is responsible for regulating transcriptional elongation in these genes. It is worth noting that most genes displayed decreased burst sizes in the PD-MK condition (Fig. 4G), suggesting that transcription elongation is not the only downstream effector of Akt and MAPK pathways regulating the normalized TBi noise.

**Fig. 6.**
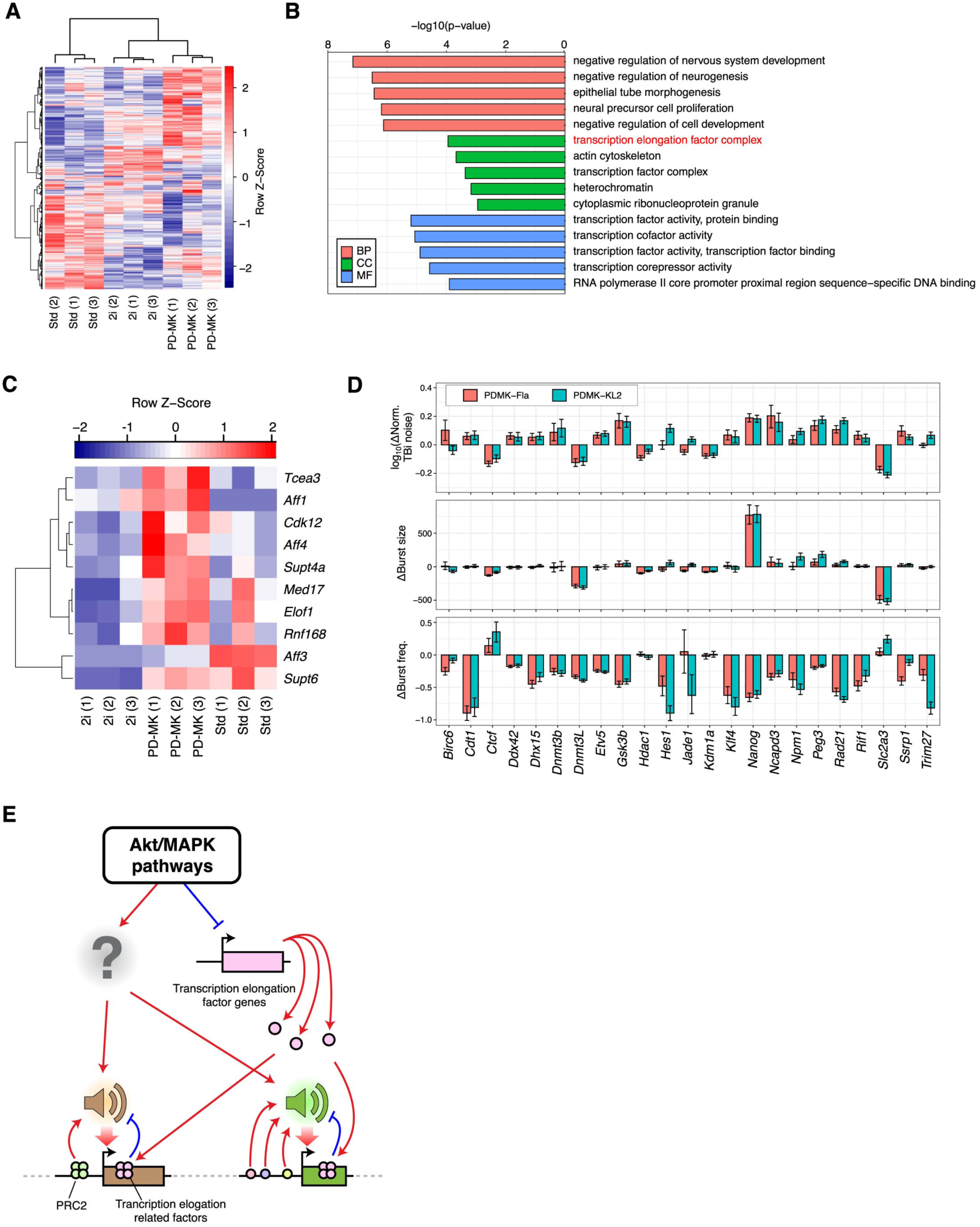
Increase in the expression level of transcription elongation factors in PD-MK condition. **(A)** Comparison of transcriptome of cells conditioned to Std, 2i, PD-MK conditions. **(B)** GO analysis of genes whose expression significantly increased in the PD-MK condition against the 2i condition. BP: biological process; CC: cellular components; MF: molecular function. **(C)** Expression levels of genes encoding transcriptional elongation factors were elevated under PD-MK condition as compared to 2i condition. **(D)** Effect of transcription elongation inhibitor Flaovopiridol and super elongation complex (SEC) inhibitor KL-2 treatment on normalized TBi noise, burst size and frequency. The KI cell lines conditioned to PD-MK were treated with Flavopiridol or KL-2 for 2 days, analyzed by flow cytometry, and normalized TBi noise, burst size and frequency were calculated. Δnormalized TBi noise, Δburst size, Δburst frequency are residuals of normalized TBi noise, burst size, frequency of inhibitor-treated from that of control cells (PD-MK condition), respectively. Error bars indicate 95% confidence interval. **(E)** Schematic summary of the determination of transcriptional bursting kinetics (including TBi noise, burst size, and burst frequency) by a combination of promoter-and gene body-binding proteins, including polycomb repressive complex 2 and transcription elongation-related factors.

## Discussion

In this study, we used scRNA-seq to measure genome-wide transcriptional bursting kinetics in hybrid mESCs. It has been reported that the promoter region of a gene is mainly involved in regulating the burst size and that enhancers are associated with the burst frequency regulation (*8*). We confirmed that the presence of the TATA box at the promoter (Fig. 2A-C) and the promoter localization of several factors (*e.g.*, EP300, ELL2, MED12, etc.) are positively correlated with the burst size (Fig. 2C). Consistent with previous reports, the enhancer localization of several factors (including transcription elongation related factors) was positively correlated with burst frequency (*8*). In addition, we found that the kinetics of transcriptional bursting was also regulated by PRC2 and transcription elongation-related factors. However, our data suggest that these individual factors are not general or master regulators of transcriptional bursting kinetics, but rather that the kinetics of individual genes are determined by a combination of promoter and gene body associated factors (Fig. 3, S5, 2F, G, 4G, 6D-E).

Not only transcriptional bursting, but also other downstream processes including post-transcriptional regulation (*22*) or translation (*32*) contribute to generating or suppressing heterogeneity in gene expression as intrinsic noise (*5*). It has been reported that TBi noise is suppressed at the protein level by nuclear retention (*22*). However, at least in 25 genes analyzed in this study in mESCs, suppression of TBi noise by nuclear retention was not observed (Fig. 1N), suggesting that noise suppression via nuclear mRNA retention is not a general phenomenon across cell types and genes.

In addition, we found a significant correlation between mRNA and protein expression levels in 25 genes examined in this study (Fig.1K). Simultaneous measurement of mRNA and protein levels in single mammalian cells showed a low correlation between them, suggesting that there could be a cause of gene expression noise even at the translation level (*32*). However, it has been reported that the average expression level of mRNA and protein significantly correlates in mammalian cells (*33*). This conflict is probably related to the difference in expression kinetics over time between mRNA and protein. That is, there is a time lag between the transcription of the mRNA and the translation of the protein, suggesting a deviation between the temporal peak of the mRNA expression amount and the temporal peak of the protein amount. However, mRNAs are transcribed from only 1 or 2 copies of the gene, which generated TBi noise during the process, whereas under typical conditions, proteins are translated from a large number of mRNAs, suggesting that the noise arising from the translation is much smaller than that from TBi noise.

Several groups have reported the detection of transcriptome-wide cell-to-cell heterogeneity in gene expression by scRNA-seq in inbred (*34, 35*) and hybrid mESC lines (*8, 36*). However, most of these studies only considered the overall heterogeneity in gene expression but not the TBi noise, resulting in incomplete characterization of the general mechanism underlying heterogeneous gene expression. Furthermore, a previous study has suggested that genes related to the cell cycle demonstrate considerable heterogeneity in expression (*36*). The phases of the cell cycle can be estimated from transcriptome data, but if the cells to be analyzed comprise a mixture of different phases of the cell cycle, the cells of a particular phase become part of the overall cells and the number of cells to be analyzed decreases. To avoid this problem, we used 447 cells in only the G1 phase for the analysis of TBi noise in our study. The larger the number of cells to be analyzed, the more reliable the calculated data regarding the kinetic properties of transcriptional bursting (Fig. S2D, E). Thus, by measuring TBi noise and heterogeneity in gene expression genome-wide, our studies enabled a detailed investigation of possible contributing factors to the regulation of transcriptional bursting and TBi noise.

Pluripotent cells in the inner cell mass share a similar transcriptomic gene expression profile with the mESCs. However, it is still poorly understood how heterogeneous gene expression could contribute to the differentiation of pluripotent cells into an epiblast and primitive endoderm (*16*). We envision future investigations to explore the involvement of TBi noise in cell fate decisions in the inner cell mass.

In conclusion, here we performed a genome-wide analysis using single-cell RNA sequencing to determine the kinetic properties of transcriptional bursting in mouse embryonic stem cells. We found that the transcriptional bursting kinetics were determined by a combination of promoter and gene body binding proteins, including transcription elongation-related factors and polycomb repressive complex 2 (PRC2). In addition, using the CRISPR lentiviral screening, we observed that the Akt/MAPK signaling pathway was also involved in this process by modulating transcription elongation efficiency. We believe that our study provides important information for understanding the molecular basis of transcriptional bursting, which underlies cellular heterogeneity.

## Material and Methods

### Cell Culture and Cell Lines

The hybrid mouse ES cell line F1-21.6 (129Sv-Cast/EiJ, female), a kind gift from Prof. Joost Gribnau, was grown on either Laminin-511 (LN511, BioLamina, Stockholm, Sweden) or gelatin coated-dish in either Std medium (15% fetal bovine serum [FBS] [Gibco], 0.1 mM β-mercaptoethanol [Wako Pure Chemicals, Osaka, Japan], and 1000U/mL of leukemia inhibitory factor [LIF, Wako Pure Chemicals, Osaka, Japan]) or 2i medium (StemSure D-MEM [Wako Pure Chemicals, Osaka, Japan], 15% of fetal bovine serum, 0.1 mM β-mercaptoethanol, 1x MEM nonessential amino acids [Wako Pure Chemicals], a 2 mM L-alanyl-L-glutamine solution [Wako Pure Chemicals], 1000 U/mL LIF [Wako Pure Chemicals], 20 mg/mL gentamicin [Wako Pure Chemicals], 1 µM PD0325901 [CS-0062, Chem Scene], and 3 µM CHIR99021 [034-23103, Wako Pure Chemicals]). This cell line was previously described in (*37*).

WT mESCs derived from inbred mouse (Bruce 4 C57BL/6J, male, EMD Millipore, Billerica, MA) and other KI derivatives were cultured on either LN511 or gelatin coated dish under either Std, 2i or PD-MK medium (StemSure D-MEM [Wako Pure Chemicals, Osaka, Japan], 15% of fetal bovine serum, 0.1 mM β-mercaptoethanol, 1x MEM nonessential amino acids [Wako Pure Chemicals], a 2 mM L-alanyl-L-glutamine solution [Wako Pure Chemicals], 1000 U/mL LIF [Wako Pure Chemicals], 20 mg/mL gentamicin [Wako Pure Chemicals], 1 µM PD0325901 [CS-0062, Chem Scene], and 4 µM MK-2206 2HCl [S1078, Selleck Chemicals, Houston TX]). Inhibitors were added at the following concentrations: 40 µM 5,6-dichloro-l-β-D-ribofuranosyl benzimidazole (DRB) (D1916, Sigma-Aldrich, St. Louis, MO); 0.25 µM Flavopiridol (CS-0018, Chem Scene, Monmouth Junction, NJ) in 2i condition; 0.125 µM Flavopiridol in PD-MK condition; 3 µM CHIR99021 (034-23103, Wako) in Std medium; 1 µM PD0325901 (CS-0062, Chem Scene) in Std medium; 5 µM BGJ398 (NVP-BGJ398) (S2183, Selleck Chemicals) in Std medium; 1 µM Rapamycin (R-5000, LC Laboratories, MA, USA) in Std medium; 0.2 µM INK128 (11811, Cayman Chemical Company, MI, USA) in Std medium; and 4 µM MK-2206 2HCl (S1078, Selleck) in Std medium.

C57BL/6NCr mESCs (male) were cultured on gelatin coated dish under PD-MK condition.

### Establishment of knock-in mESC lines

To quantify the TBi noise level of a particular gene, it is necessary to establish a cell line with the GFP and iRFP reporter genes individually knocked into both alleles of the target genes. Therefore, based on the scRNA-seq data, 25 genes (Fig 1H) with medium expression levels and variable TBi noise levels were manually selected.

GFP/iRFP knock-in cell lines were established using CRISPR-Cas9 or TALEN expression vectors and targeting vectors (with about 1-kbp homology arms). Vectors used in this study are listed in Table S4. C57BL/6J mESCs (5 × 10^5^) conditioned to 2i medium were plated onto gelatin-coated 6-well plates. After 1 hour, the cells were then transfected with 1 µg each of GFP and iRFP targeting vectors (Table S4), 1 µg total of nuclease vectors (Table S4), and pKLV-PGKpuro2ABFP (puromycin resistant, Addgene, Plasmid #122372) using Lipofectamine 3000 (Cat# L3000015, Life Technologies, Gaithersburg, MD), according to the manufacturer’s instructions. Cells were selected by adding puromycin (1 µg/mL) to the 2i medium 24 h post-transfection. After another 24 hours the medium was exchanged. The medium was exchanged every two days. At 5 days after transfection, cells were treated with 25 µM biliverdin (BV). BV is used for forming a fluorophore by iRFP670. Although BV is a molecule ubiquitous in eukaryotes, the addition of BV to culture medium increases the fluorescence intensity. Twenty-four hours later, cells were trypsinized, subjected to FACS analysis, GFP/iRFP double-positive cells were sorted, and seeded on a gelatin-coated 6-cm dish. The medium was exchanged every two days. One week after sorting, 16 colonies were picked for downstream analysis and checking gene targeting. PCR was carried out using primers outside the homology arms, and cells that seemed to be successfully knocked into both alleles were selected. Thereafter, candidate clones were further analyzed by Southern blotting as described before (*18*)(Fig. S3). Restriction enzymes and genomic regions used for Southern blot probes are listed in Table S4. Probes were prepared using the PCR DIG Probe Synthesis Kit (Roche Diagnostics, Mannheim, Germany).

### Mice

ICR mice were purchased from CLEA Japan (Tokyo, Japan). All mice were housed in an air-conditioned animal room under specific-pathogen-free (SPF) conditions, with a 12 h light/dark cycle. All mice were fed a standard rodent CE-2 diet (CLEA Japan, Tokyo, Japan) and had *ad libitum* access to water. All animal experiments were approved by the President of NCGM, following consideration by the Institutional Animal Care and Use Committee of NCGM (Approval ID: No. 18037), and were conducted in accordance with institutional procedures, national guidelines and relevant national laws on the protection of animals.

### Plasmid construction

To construct the lentiCRISPRv2-sgSuz12_1, lentiCRISPRv2-sgSuz12_2, lentiCRISPRv2-sgSuz12_3, and lentiCRISPRv2_sgMS2_1 plasmids, which are sgRNA expression vectors, we performed inverse PCR using R primer (5’-GGTGTTTCGTCCTTTCCACAAGAT-3’) and either of F primers (5’-AAAGGACGAAACACCGCGGCTTCGGGGGTTCGGCGGGTTTTAGAGCTAGAAATAGCAAGT-3’, 5’-AAAGGACGAAACACCGGCCGGTGAAGAAGCCGAAAAGTTTTAGAGCTAGAAATAGCA AGT-3’, 5’-AAAGGACGAAACACCGCATTTGCAACTTACATTTACGTTTTAGAGCTAGAAATAGCAA GT-3’, or 5’-AAAGGACGAAACACCGGGCTGATGCTCGTGCTTTCTGTTTTAGAGCTAGAAATAGCAAGT-3’), respectively, and lentiCRISPR v2 (Addgene, Plasmid #52961) as a template, followed by self-circularization using the In-Fusion HD Cloning Kit (Cat# 639648, Clontech Laboratories, Mountain View, CA, USA).

### RNA-Seq of mESCs cultured on LN511 and serum-LIF medium

129/CAST hybrid mESCs need to be maintained on feeder cells in gelatin/Std condition. To eliminate the need for feeder cells, we decided to maintain the hybrid mESCs on dishes coated with LN511 enabling maintenance of mESCs without feeder cells in Std condition. To compare the transcriptomes of mESCs cultured on gelatin-coated dish and those cultured on Laminin-511 coated dish, we performed RNA-Seq analysis as follows. First, C57BL/6J WT mESCs were conditioned on either gelatin or LN511 coated dish in either Std or 2i medium for 2 weeks. Next, RNA was recovered from 1 x 10^6^ cells using the NucleoSpin RNA kit (Macherey-Nagel, Düren, Germany). The RNA was sent to Eurofins for RNA-seq analysis. RNA-seq reads were aligned to the mouse reference genome (mm10) using TopHat (version 2.1.1) (https://ccb.jhu.edu/software/tophat/index.shtml). Fragments per kilobase per million mapped reads (FPKM) values were quantified using Cufflinks (version 2.1.1) (http://cole-trapnell-lab.github.io/cufflinks/) to generate relative gene expression levels. Hierarchical clustering analyses were performed on FPKM values using CummeRbund (v2.18.0) (https://bioconductor.org/packages/release/bioc/html/cummeRbund.html). On comparison, the transcriptomes of mESCs cultured on gelatin-coated dish and those cultured on Laminin-511 coated dishes showed no considerable difference in expression patterns (Fig. S1A).

### Sequencing library preparation for RamDA-seq

Library preparation for single-cell RamDA-seq was performed as described previously (*20*). Briefly, hybrid mESC line F1-21.6 (129Sv-Cast/EiJ) conditioned to LN511/Std condition were dissociated with 1× Trypsin (Thermo Fisher Scientific, Rochester, NY) with 1 mM EDTA at 37 °C for 3 min. The dissociated cells were adjusted to 1 × 10^6^ cells/mL and stained with 10 μg/mL Hoechst 33342 dye (Sigma-Aldrich) in phosphate-buffered saline (PBS) at 37 °C for 15 min to identify the cell cycle. After Hoechst 33342 staining, the cells were washed once with PBS and stained with 1 μg/mL propidium iodide (PI, Sigma-Aldrich) to remove dead cells. Single-cell sorting was performed using MoFlo Astrios (Beckman Coulter). Recent studies of scRNA-Seq using mESCs have suggested that genes related to the cell cycle demonstrate considerable heterogeneity in expression (*36*). Therefore, in order to minimize this variation, 474 cells only in the G1 phase were collected. Single cells were collected in 1 μL of cell lysis buffer (1 U RNasein plus [Promega, Madison, WI], 10% RealTime ready Cell Lysis Buffer [Cat# 06366821001, Roche], 0.3% NP40 (Thermo Fisher Scientific), and RNase-free water (TaKaRa, Japan)) in a 96-well PCR plate (BIOplastics).

The cell lysates were denatured at 70 °C for 90 s and held at 4 °C until the next step. To eliminate genomic DNA contamination, 1 µL of genomic DNA digestion mix (0.5× PrimeScript Buffer, 0.2 U of DNase I Amplification Grade, 1: 5 000 000 ERCC RNA Spike-In Mix I (Thermo Fisher Scientific) in RNase-free water) was added to 1 µL of the denatured sample. The mixtures were agitated for 30 s at 2000 rpm using a ThermoMixer C at 4 °C, incubated in a C1000 thermal cycler at 30 °C for 5 min and held at 4 °C until the next step. One microliter of RT-RamDA mix (2.5× PrimeScript Buffer, 0.6 pmol oligo(dT)18 (Cat# SO131, Thermo Fisher Scientific), 8 pmol 1st-NSRs (*20*), 100 ng of T4 gene 32 protein (New England Biolabs), and 3× PrimeScript enzyme mix (Cat# RR037A, TAKARA Bio INC) in RNase-free water) was added to 2 µL of the digested lysates. The mixtures were agitated for 30 s at 2,000 rpm and 4 °C, and incubated at 25 °C for 10 min, 30 °C for 10 min, 37 °C for 30 min, 50 °C for 5 min, and 94 °C for 5 min. Then, the mixtures were held at 4 °C until the next step. After RT, the samples were added to 2 µL of second-strand synthesis mix (2.5× NEB buffer 2 [New England Biolabs], 0.625 mM each dNTP Mixture [TaKaRa], 40 pmol 2nd-NSRs (*20*), and 0.75 U of Klenow Fragment [3’ → 5’ exo-] [New England Biolabs] in RNase-free water). The mixtures were agitated for 30 s at 2,000 rpm and 4 °C, and incubated at 16°C for 60 min, 70°C for 10 min and then at 4 °C until the next step. Sequencing library DNA preparation was performed using the Tn5 tagmentation-based method with 1/4 volumes of the Nextera XT DNA Library Preparation Kit (Cat# FC-131-1096, −2001,−2002, −2003, and −2004, Illumina, San Diego, CA) according to the manufacturer’s protocol. The above-described double-stranded cDNAs were purified by using 15 μL of AMPure XP SPRI beads (Cat# A63881, Beckman Coulter) and a handmade 96-well magnetic stand for low volumes. Washed AMPure XP beads attached to double-stranded cDNAs were directly eluted using 3.75 μL of 1× diluted Tagment DNA Buffer (Illumina) and mixed well using a vortex mixer and pipetting. Fourteen cycles of PCR were applied for the library DNA. After PCR, sequencing library DNA was purified using 1.2× the volume of AMPure XP beads and eluted into 24 μL of TE buffer.

### Quality control and sequencing of library DNA

All the RamDA-seq libraries prepared with Nextera XT DNA Library Preparation were quantified and evaluated using a MultiNA DNA-12000 kit (Shimadzu, Kyoto, Japan) with a modified sample mixing ratio (1:1:1; sample, marker, and nuclease-free water) in a total of 6 μL. The length and yield of the library DNA were calculated in the range of 161–2,500 bp. The library DNA yield was estimated as 0.5 times the value quantified from the modified MultiNA condition. Subsequently, we pooled each 110 fmol of library DNA in each well of a 96-well plate. The pooled library DNA was evaluated based on the averaged length and concentration using a Bioanalyzer Agilent High-Sensitivity DNA Kit (Cat# 5067-4626) in the range of 150–3,000 bp and a KAPA library quantification kit (Cat# KK4824, Kapa Biosystems, Wilmington, MA). Finally, 1.5 pM pooled library DNA was sequenced using Illumina HiSeq2000 (single-read 50 cycle sequencing).

### smFISH

2×10^5^ trypsinized cells were transferred onto LN511 coated round coverslips and cultured for 1 h at 37°C and 5% CO_2_. Cells were washed with PBS, fixed with 4% paraformaldehyde in PBS for 10 min, and washed with PBS two times. Then, cells were permeabilized in 70% ethanol at 4°C overnight. Following a wash with 10% formamide dissolved in 2× SSC, the cells were hybridized to probe sets in 60 μL of hybridization buffer containing 2× SSC, 10% dextran sulfate, 10% formamide, and each probe set (Table S4). Hybridization was performed for 4 h at 37°C in a moist chamber. The coverslips were washed with 10% formamide in 2× SSC solution, and then with 10% formamide in 2× SSC solution with Hoechst 33342 (1:1000). Hybridized cells were mounted in catalase/glucose oxidase containing mounting media [0.4% glucose in 10 mM Tris, 2X SSC, 37 µg/ml glucose oxidase, 1/100 catalase (Sigma-Aldrich, C3155)]. Images were acquired using a Nikon Ti-2 microscope with a CSU-W1 confocal unit, a 100× Nikon oil-immersion objective of 1.49 NA, and an iXon Ultra EMCCD camera (Andor, Belfast, UK), with laser illumination at 405 nm, 561 nm, and 637 nm, and were analyzed using NIS-elements software (version 5.11.01, Nikon, Tokyo, Japan); 101 z planes per site spanning 15 μm were acquired. Images were filtered with a one-pixel diameter 3D median filter and subjected to background subtraction via a rolling ball radius of 5 pixels, using FIJI software. Detection and counting of smFISH signals were performed using FISHquant software version 3 (https://bitbucket.org/muellerflorian/fish_quant/src/master/).

FISHquant quantifies the number of mRNAs in the cell nucleus and cytoplasm. Mixtures of mNeonGreen and iRFP670 probes conjugated with CAL Fluor Red 590 and Quasar 670 were obtained from BioSearch Inc (Novato, CA) and used at 0.25 µM. Probe sequences are shown in Table S4. TBi noise was calculated in the same way as described in “Analysis of scRamDA-seq data for individual transcripts” section. Since smFISH has almost the same average value, correction between alleles was not carried out. The count normalized log-ratios of TBi noise (normalized TBi noise) were calculated as the residuals of the regression line (Fig. S1H). Normalization by gene length had not been applied for the smFISH data.

### Flow cytometry analysis for calculation of TBi noise

On the day before flow cytometry, cells were treated with 25 µM BV. Cells that became 80% confluent were washed with PBS, trypsinized, inactivated with FluoroBrite DMEM (Thermo Fisher Scientific) containing 10% FBS, and centrifuged to collect the cells. Cells were suspended in PBS to be 1 to 5 × 10^6^ cells/mL. Fluorescence data of SSC, FSC, GFP and iRFP were obtained with BD FACS Aria III. Cells were gated based on FSC and SSC using a linear scale to gate out cellular debris. Among GFP and iRFP data, extreme values indicating 20 * interquartile range (IQR) or more were excluded from analysis. The mean value of the negative control data of WT mESC was subtracted from the data to be analyzed, and the data that fell below zero were excluded. Furthermore, correction was made using the following equation so that the mean fluorescent intensities between GFP and iRFP were consistent.

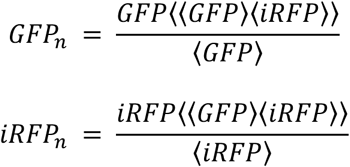

Here, the *i* th element of vectors *GFP* and *iRFP* contains the fluorescent intensities of GFP and iRFP, respectively, of the *i* th cell in the sample. *GFP_n_* and *iRFP_n_* represent mean normalized *GFP* and *iRFP*, respectively. Then, TBi noise is calculated as described in “Analysis of scRamDA-seq data for individual transcripts” section. The relationship between mean fluorescent intensities and TBi noise was plotted (Fig. S1I). The fluorescent intensity normalized log-ratios of TBi noise (normalized TBi noise) were calculated as the residuals of the regression line (Fig. S1I).

### Immunofluorescence

On the day before immunostaining, *Trim28*, *Dnmt3l*, *Klf4*, *Peg3*, *Npm1*, *Dnmt3b*, *Nanog*, *Rad21*, and *Hdac1* KI cell lines at ∼ 70% confluence were treated with 25 μM BV. After 24 hours, 1 × 10^5^ cells were plated onto the 8-well Lab-Tek II chambered coverglass (Thermo Fisher Scientific) coated with LN511. For immunostaining of C57B6J WT mESCs conditioned to Std/LN511, 2i/LN511, and PD-MK/LN511 condition, cells were plated 1 x 10^5^ onto an 8-well Lab-Tek II chambered coverglass coated with LN511. After 1 hour, cells were washed once with PBS and fixed with 4% PFA for 10 min at room temperature. Fixed cells were washed with BBS buffer (50 mM BES, 280 mM NaCl, 1.5 mM Na_2_HPO_4_・2H_2_O, and 1 mM CaCl_2_) two times, and blocked for 30 min in BBT-BSA buffer (BBS with 0.5% BSA, 0.1% Triton, and 1 mM CaCl_2_) at room temperature. Cells with primary antibodies were incubated overnight at 4°C at the following dilutions: anti-TRIM28 (1:500; GTX102227, GeneTex, RRID:AB_2037323), anti-DNMT3L (1:250; ab194094, Abcam, Cambridge, MA, RRID:AB_2783649), anti-KLF4 (1:250, ab151733, Abcam, RRID:AB_2721027), anti-PEG3 (1:500; BS-1870R, Bioss Antibodies, RRID:AB_10855800), anti-NPM1 (1:100; A302-402A, Bethyl Labolatories Inc., RRID:AB_1907285), anti-DNMT3B (1:500; 39207, Active motif, RRID:AB_2783650), anti-NANOG (1:500, 14-5761-80, eBioscience, RRID:AB_763613), anto-RAD21 (1:500; GTX106012, GeneTex, RRID:AB_763613), anti-HDAC1 (1:500; GTX100513, GeneTex, RRID:AB_1240929), anti-OCT-4A (1:400; 2840, Cell Signaling Technology, RRID:AB_2167691), and anti-SSEA1 (1:1000; 4744, Cell Signaling Technology, RRID:AB_1264258). Cells were washed and blocked in BBT-BSA. Then, for KI cell lines, cells were incubated with Alexa Fluor 594 conjugated secondary antibodies (1:500, Life Technologies). For C57B6J WT mESCs, cells were incubated with Alexa Fluor 488 goat anti-mouse IgG, Alexa Fluor 594 goat anti-rabbit IgG, and Alexa Fluor 647 goat anti-rat IgG secondary antibodies (1:500, Life Technologies). Images were acquired using a Nikon Ti-2 microscope with a CSU-W1 confocal unit, a 100× Nikon oil-immersion objective of 1.49 NA, and an iXon Ultra EMCCD camera (Andor, Belfast, UK).

### Suz12 knockout

*Dnmt3l*, *Dnmt3b*, *Peg3*, and *Ctcf* KI cell lines conditioned to gelatin/2i condition were trypsinized, and plated onto a 24-well plate at 5 × 10^5^ cells/500 μL each. One hour later, for *Suz12* knockout (KO), 330 ng each of lentiCRISPRv2-sgSuz12_1, lentiCRISPRv2-sgSuz12_2, lentiCRISPRv2-sgSuz12_3, and 300 ng of pCAG-mTagBFP2 (Addgene, Plasmid #122373) plasmids, or for control, 1000 ng of lentiCRISPRv2_sgMS2_1 and 300 ng pCAG-mTagBFP2 (Addgene, Plasmid #122373) plasmids were transfected using Lipofectamine 3000 into each cell line. Two days later, BFP positive cells were sorted by FACS, and plated onto a 6-cm dish. After 1 week, we picked up 8 colonies for *Suz12* KO, and 4 colonies for control for downstream analysis. We checked the expression of PRC2 related proteins by western blotting (see below). Then, cells were conditioned to LN511/Std medium for at least 2 weeks. As described above, flow cytometry analysis was performed to calculate normalized TBi noise, burst size, and burst frequency.

### Western blotting

Cells are washed twice with PBS, trypsinized and collected by centrifugation. Cells were counted and then washed twice with PBS. Finally, cells were lysed in the lysis buffer (0.5% Triton X-100, 150 mM NaCl, 20 mM Tris-HCl, pH 7.5) to obtain 1 × 10^6^ cells/100 μL. Then, the lysates were incubated at 95℃ for 5 min, and filtered by QIAshredder homogenizer (QIAGEN). The extracted proteins were analyzed by 5-20% gradient SDS-PAGE and transferred onto Immobilon transfer membranes (Millipore, Billerica, MA, USA) for immunoblotting analyses. The primary antibodies used were anti-SUZ12 (1:1000; 3737, Cell Signaling Technology, RRID:AB_2196850), anti-EZH2 (1:1000; 5246, Cell Signaling Technology, RRID:AB_10694683), anti-Histone H3K27me3 (1:1000; 39155, Active Motif, RRID:AB_2561020), anti-GAPDH (1:5000; 5174, Cell Signaling Technology, RRID:AB_10622025), anti-Phospho-MEK1/2 (Ser217/221) (1:1000; 9154, Cell Signaling Technology, RRID:AB_2138017), anti-MEK1/2 (1:1000; 8727, Cell Signaling Technology, RRID:AB_10829473), anti-p44/42 MAPK (Erk1/2) (1:1000; 4695, Cell Signaling Technology, RRID:AB_390779), anti-Phospho-p44/42 MAPK (Erk1/2) (Thr202/Tyr204) (1:2000; 4370, Cell Signaling Technology, RRID:AB_2315112), anti-Phospho-4E-BP1 (Thr37/46) (1:1000; 2855, Cell Signaling Technology, RRID:AB_560835), anti-Phospho-Akt (Ser473) (1:1000; 4060, Cell Signaling Technology, RRID:AB_2315049), anti-Phospho-Akt (Thr308) (1:1000; 13038, Cell Signaling Technology, RRID:AB_2629447), anti-Akt (pan) (1:1000; 4691, Cell Signaling Technology, RRID:AB_915783), anti-c-Myc (1:1000; ab32072, Abcam, RRID:AB_731658), anti-FoxO1 (1:1000; 14952, Cell Signaling Technology, RRID:AB_2722487), anti-FOXO3A (1:2500; ab12162, Abcam, RRID:AB_298893), anti-Nanog (1:500, 14-5761-80, eBioscience, RRID:AB_763613), anti-OCT-4A (1:500; 2840, Cell Signaling Technology, RRID:AB_2167691), and anti-SOX2 (1:1000; ab97959, Abcam, RRID:AB_2341193).

### Infection of CRISPR lentivirus library

*Nanog*, *Trim28*, and *Dnmt3L* KI cells were transduced with the Mouse CRISPR Knockout Pooled Library (GeCKO v2) (Addgene, # 1000000052) (*29*) via spinfection as previously described. We used only Mouse library A gRNA. Briefly, 3 x 10^6^ cells per well (a total of 1.2 x 10^7^ cells) were plated into a LN511-coated 12 well plate in the standard media supplemented with 8 µg/mL polybrene (Sigma-Aldrich). Each well received a virus amount equal to MOI = 0.3. The 12-well plate was centrifuged at 1,000 g for 2 h at 37°C. After the spin, media were aspirated and fresh media (without polybrene) was added. Cells were incubated overnight. Twenty-four hours after spinfection, cells were detached with trypsin and were replated into 4 of LN511-coated 10 cm dishes with 0.5 µg/mL puromycin for 3 days. Media were refreshed daily. At 6 days after transduction, cells were treated with 25 µM BV. After 24 h, at least 1.75 × 10^5^ cells showing GFP/iRFP expression ratio close to 1 were sorted by FACS and plated on 12 well plates (LM 511/Std condition). Unsorted cells were passaged to 10 cm plates, 5 × 10^5^ each. After the expansion of these sorted cells for 1 week, cells with GFP/iRFP expression ratio were close to 1 were sorted again. These sorting and expansion procedures were repeated 4 times in total. After 3 days after the fourth sorting, 2 x 10^5^ cells were collected and genomic DNA was extracted. PCR of the virally integrated guides was performed on sgDNA at the equivalent of approximately 2000 cells per guide in 48 parallel reactions using KOD-FX neo (TOYOBO, Japan) in a single-step reaction of 22 cycles. Primers are listed here: forward primer, AATGATACGGCGACCACCGAGATCTACACTCTTTC CCTACACGACGCTCTTCCGATCTNNNNNNNN(1–8-bp stagger) GTGGAAAGGACGAAACACCG; reverse primer, CAAGCAGAAGACGGCATACGAGAT*NNNNNNNN* GTGACTGGAGTTCAGACGTGTGCTCTTCCGATCTTGTGGGCGATGTGCGCTCTG, 8-bp index read barcode indicated in italics. PCR products from all 48 reactions were pooled, purified using PCR purification kit (Qiagen, Hilden, Germany) and gel extracted using the Gel extraction kit (Qiagen, Hilden, Germany). The resulting libraries were deep-sequenced on Illumina HiSeq platform with a total coverage of >8 million reads passing filter per library.

### Cell cycle analysis

4 × 10^5^ cells were seeded on LN511-coated 6-well plates. After overnight culture, the cells were incubated for 1 h with 5-ethynyl-2-deoxyuridine (EdU) diluted to 10 μM in the indicated ES cell media. All samples were processed according to the manufacturer’s instructions (Click-iT™ Plus EdU Alexa Fluor™ 647 Flow Cytometry Assay Kit, Cat# C10634, Thermo Fisher Scientific). EdU incorporation was detected by Click-iT chemistry with an azide-modified Alexa Fluor 647. Cells were resuspended in EdU permeabilization / wash reagent and incubated for 30 min with Vybrant DyeCycle Violet (Thermo Fisher Scientific). Flow cytometric was performed on a FACS Aria III (BD) and analyzed with Cytobank (www.cytobank.org) (Cytobank Inc, Santa Clara, CA).

### Analysis of apoptosis

Annexin V staining was performed using Annexin V Apoptosis Detection Kit APC (Cat# 88-8007-72, Thermo Fisher Scientific) as described in the manufacturer’s manual. Briefly, cells were trypsinized and centrifuged, and then the supernatant was removed. The remaining cells were resuspended in PBS and counted. Cells were washed once with PBS, and then resuspended in 1x Annexin V binding buffer at 1-5 x 10^6^ cells/mL. Pellets were resuspended in 100 μL of Annexin V buffer to which 5 μL of fluorochrome-conjugated Annexin V was added. Cells were incubated in the dark at RT for 15 min, washed in 1X Binding Buffer and resuspend in 200 μL of 1X Binding Buffer. Add 5 μL of Propidium Iodide Staining Solution and immediately analyzed by flow cytometry.

### Bulk RNA-Seq

RNA was extracted from either Std/LN511, 2i/LN511 or PDMK/LN511 conditioned cells at 70% confluency in a well of a 6-well plate using RNeasy Plus Mini (Qiagen). Three biological replicates were prepared. Bulk RNA-Seq was performed by CEL-Seq2 method (*38*) with total RNA amounts were used the range of 30-60 ng. The resulting reads were aligned to the reference genome (GRCm38) using HISAT (v.2.1.0) (https://ccb.jhu.edu/software/hisat/index.shtml). The software HTSeq (version 0.6.1) (https://htseq.readthedocs.io/en/release_0.11.1) was used in calculating gene-wise unique molecular identifier (UMI) counts that were converted into transcript counts after collision probability correction (*39*). The counts were input to the R library DESeq2 (version 1.14.1) (https://bioconductor.org/packages/release/bioc/html/DESeq2.html) for DE analysis. The genes that increased significantly (adjusted *P* <0.05) in the PD-MK condition against the 2i condition were subjected to GO analysis using an R package, clusterProfiler (v3.9.2) (https://bioconductor.org/packages/release/bioc/html/clusterProfiler.html).

### RNA degradation rate determination using 4sU pulse labeling

C57BL6/J WT mESCs conditioned to LN511/Std or LN511/PD-MK conditions were treated with 400 µM 4-thiouridine (4sU) for 20 min. Then, RNA was extracted from more than 1 × 10^7^ cells using RNeasy Plus Mini Kit (Qiagen, Valencia, CA). Three biological replicates were prepared for each condition. We synthesized mRuby2 RNA for spike-in RNA by standard PCR, *in vitro* transcription using T7 High Yield RNA Synthesis Kit (Cat# E2040, New England Biolabs) and purification with RNeasy Plus Mini Kit (Qiagen, Valencia, CA). Biotinylation of 4sU-labeled RNA was carried out in RNase-free water with 10 mM Tris-HCl (pH 7.4), 1 mM EDTA and 0.2 mg/mL Biotin-HPDP at a final RNA concentration of 1 μg/μL extracted RNA (a total of 125 µg) with 125 ng/µL of spike-in RNA for 3 h in the dark at room temperature. To purify biotinylated RNA from an excess of Biotin-HPDP, a Phenol:Chloroform:Isoamylalcohol (v/v = 25:24:1, Nacalai Tesque, Kyoto, Japan) extraction was performed. Phenol: Chloroform: Isoamylalcohol was added to the reaction mixture in a 1:1 ratio, followed by vigorous mixing, and centrifuged at 20,000 × g for 5 min at 4°C. The RNA containing aqueous phase was removed and transferred to a fresh, RNase-free tube. To precipitate RNA, 1/10 reaction volume of 5 M NaCl and an equal volume of 2-propanol was added and incubated for 10 min at room temperature. Precipitated RNA was collected through centrifugation at 20,000 *g* for 30 min at 4°C. The pellet was washed with an equal volume of 75% ethanol and precipitated again at 20,000 *g* for 20 min. Finally, RNA was reconstituted in 25-50 µL of RNase-free water. For removing of biotinylated 4sU-RNA, streptavidin-coated magnetic beads (Dynabeads MyOne Streptavidin C1 beads, ThermoFisher) were used according to the manufacturer’s manual. To avoid unfavorable secondary RNA structures that potentially impair the binding to the beads, the RNA was first denatured at 65°C for 10 min followed by rapid cooling on ice for 5 min. 200 µL of Dynabeads magnetic beads per sample was transferred to a new tube. An equal volume of 1× B&W (5 mM Tris-HCl, pH7.5, 0.5 mM EDTA, 1M NaCl) was added to the tube and mixed well. The tube was placed on a magnet for 1 min and the supernatant was discarded. The tube was removed from the magnet. The washed magnetic beads were resuspended in 200 µL of 1× B&W. The bead washing step was repeated for a total of 3 times. The beads were washed twice in 200 µL of Solution A (DEPC-treated 0.1 M NaOH, DEPC-treated 0.05 M NaCl) for 2 min. Then, the beads were washed once in 200 µL of Solution B (DEPC-treated 0.1 M NaCl). Washed beads were resuspended in 400 µL of 2× B&W Buffer. An equal volume of 20 µg of biotinylated RNA in distilled water was added. The mixture was incubated for 15 min at room temperature with gentle rotation. The biotinylated RNA coated beads were separated with a magnet for 2–3 min. Unbound (unbiotinylated) RNA from the flow-through was recovered using the RNeasy MinElute kit (Qiagen) and reconstituted in 25 μL of RNase-free water. cDNA was synthesized with the ReverTra Ace qPCR RT kit (Cat# FSQ-101, TOYOBO, Japan) from both total RNA and unbound (unbiotinylated) RNA. The relative amount of existing RNA (unbiotinylated RNA) / total RNA was quantified by qPCR with THUNDERBIRD SYBR qPCR Mix (Cat# QPS-201, TOYOBO). cDNAs derived from total and unbound RNA, and primers used are listed in Table S4.

### Generation of chimaeras

C57BL/6NCr ES cells derived from C57BL/6NCr (Japan SLC, Hamamatsu, Japan) were cultured in PD-MK medium on a gelatin coated dish for 2 weeks. The day before injection, the culture medium was changed to standard medium. mESCs were microinjected into eight-cell stage embryos from ICR strain (CLEA Japan, Tokyo, Japan). The injected embryos were then transferred to the uterine horns of appropriately timed pseudopregnant ICR mice. Chimeras were determined by the presence of black eyes at birth, and by coat color around 10 days after birth.

### Quantification of gene and allelic expression level

For each scRamDA-seq library, the FASTQ files of sequencing data with 10 pg of RNA were combined. Fastq-mcf (version 1.04.807) (https://github.com/ExpressionAnalysis/ea-utils/blob/wiki/FastqMcf.md) was used to trim adapter sequences and generate read lengths of 50 nucleotides (nt) with the parameters “-L 42 -l 42 -k 4 -q 30 -S.”. The reads were mapped to the mouse genome (mm10) using HISAT2 (version 2.0.4) (https://ccb.jhu.edu/software/hisat2/index.shtml) with default parameters. We removed 27 abnormal samples showing abnormal gene body coverage of sequencing reads by human curation. Using the remaining data derived from 447 cells, allelic gene expressions were quantified using EMASE (version 0.10.11) with default parameters (https://github.com/churchill-lab/emase). 129 and CAST genomes by incorporating SNPs and indels into reference genome and transcriptome was created by Seqnature (https://github.com/jaxcs/Seqnature). Bowtie (version 1.1.2) (http://bowtie-bio.sourceforge.net/index.shtml) was used to align scRamDA-seq reads against the diploid transcriptome with the default parameters.

### Estimation of the kinetic properties of transcriptional bursting using transcript-level count data

To calculate TBi noise, it is assumed that the average expression levels among the alleles are equal. For this purpose, firstly global allelic bias in expression level was normalized using TMM normalization method implemented in the R package edgeR (https://bioconductor.org/packages/release/bioc/html/edgeR.html). The total noise 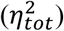 for each TC was calculated using the following equation (4).

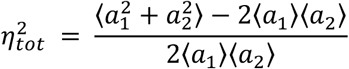

Here, the *i* th element of vectors *a_1_* and *a_2_* contains the read counts of TC from allele 1 or allele 2, respectively, of the *i* th cell in the sample. Global normalization does not substantially change the shape of the read count-total noise distribution (Fig. S1B). Then, the read counts were normalized between alleles at each transcript by the normalize.quantiles.robust method using the Bioconductor preprocessCore package (version 1.38.1)

(https://bioconductor.org/packages/release/bioc/html/preprocessCore.html). Furthermore, correction was made using the following equation so that the mean read counts among the alleles were consistent.

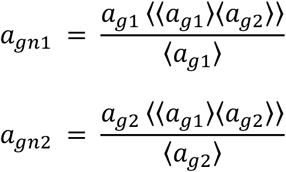

Here, the *i* th element of vectors *a_g1_* and *a_g2_* contains the globally and allelically normalized read counts of TC from allele 1 or allele 2, respectively, of the *i* th cell in the sample. *a_gn1_* and *a_gn2_* represent mean normalized *a_g1_* and *a_g2_*, respectively. Angled brackets denote means over the cell population. From these read count matrices, the TBi noise 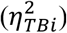 for each TC was calculated using the following equation (*4*).

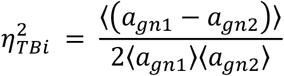

TCs showing the relatively large difference in expression level between alleles before correction (the average pre-normalized expression level between alleles was more than 100 read counts) was excluded from the subsequent analysis. We found that some TCs showed TBi noise below Poisson noise (Fig. S1C). Theoretically, TBi noise cannot be below Poisson noise (*12*), suggesting that these have some possibility of analytical inadequacies, and were excluded from the downstream analysis. A decrease in TBi noise was observed as the expression level increased as theoretically expected (Fig. S1C). To investigate the factors involved in the TBi noise and bursting properties independent of expression level, the count normalized log-ratios of TBi noise were calculated as the residuals of a regression line that is calculated using data set with more than 1 mean read counts (Fig. S1D). In addition, global correlation was found between the length of the transcript and the count normalized TBi noise (Fig. S1E). Thus, the count and transcript length normalized log-ratios of TBi noise were calculated as the residuals of a regression line (Fig. S1E, F). We call these read count and transcript length normalized TBi noise simply normalized TBi noise. For transcripts with low expression levels, it is difficult to distinguish whether their heterogeneity in expression level is due to technical or biological noise. Therefore, transcripts with read count less than 20 were excluded from the downstream analysis (remaining 5,992 TCs).

TBi noise is a function of mRNA degradation rate (Fig. 1A) (*7, 9, 11*). The mRNA degradation rate in mESC has been genome-wide analyzed (*21*). Genes whose degradation rate is unknown were provisionally assigned a median value. The burst size (*b*) and burst frequency (*f*) of each transcript can be estimated by mRNA degradation rate (*γ_m_*), TBi noise 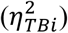 and mean number of mRNA (*µ*) as the following equations (*7, 9, 11*).

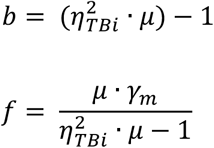

Previous studies have reported the estimation of the burst size and burst frequency for each allele using a Poisson-Beta hierarchical model with scRNA-seq data of hybrid cells (*8, 40*). To evaluate the validity of the parameters derived from the above mentioned equations, we used our hybrid mESC scRNA-seq data and the SCALE software (version 1.3.0) that enables to estimate the burst frequency and burst size per allele using a Poisson-Beta hierarchical model (*40*). Since the SCALE software always sets the RNA degradation rate to 1, the resulting parameters can be considered as RNA degradation-rate normalized parameters. Therefore, for comparison, the burst frequency calculated from TBi noise was divided by the RNA degradation rate to obtain RNA degradation-rate normalized burst frequency (see above formula). We found that the SCALE-based and TBi-noise-based parameters were well correlated (R > 0.8) (Fig S2F-K), suggesting that the burst size and burst frequency calculated using TBi noise are valid.

### Estimation of the kinetic properties of transcriptional bursting using gene-level count data

It is thought that the RNA detected by smFISH is not a specific TC but contains multiple transcript variants. Therefore, TBi noise data calculated using TC-level count data could not be compared to that from smFISH data. To solve this problem, scRamDA-seq data for each TC were summed up for each gene, and TBi noise was recalculated. For this purpose, global allelic bias in expression level was first normalized as described above. Then, data of each TC were summed up for each gene at this time point. Next, the read counts were normalized between alleles at each gene by the normalize.quantiles.robust method using the Bioconductor preprocessCore package. Furthermore, correction was made so that the mean read counts among the alleles were consistent as described above. From these read count matrices, the TBi noise for each gene can be calculated as described above. Data with TBi noise below Poisson noise were excluded from the downstream analysis. To investigate the factors involved in the TBi noise and bursting properties independent of expression level, the count normalized log-ratios of TBi noise were calculated as the residual of a regression line that is calculated using a data set with more than 1 mean read counts. Then, the count and gene length normalized log-ratios of TBi noise were calculated as the residual of a regression line. We call these read count and gene length normalized TBi noise simply normalized TBi noise. The burst size (*b*) and burst frequency (*f*) of each gene can be estimated as described above.

### TATA box identification

We used FindM (https://ccg.epfl.ch/ssa/findm.php) to determine whether a sequence of 50 bp upstream from the TSS of transcripts, with an average read count of over 20 in our scRNA-seq, contained a TATA box.

### Correlation analysis

We used bioinformatics tools freely available on Galaxy Project platform (https://galaxyproject.org/). Various ChIP-seq data were obtained from the bank listed in Table S4. Then, we mapped them to mm10 genome with Bowtie (Galaxy Version 1.1.2), converted it to bam file with SAM-to-BAM tool (Galaxy Version 2.1). Reads Per Million mapped reads (RPM) data from −1,000 to +100 from TSS, and gene body of individual transcripts were analyzed by the ngs.plot (version 2.61) (https://github.com/shenlab-sinai/ngsplot). Of these, extreme outliers (100 times the average value) were excluded from analysis. In addition, we also considered the replication timing, promoter proximal pausing of RNA polymerase II, considered to be related to the characteristics of transcriptional bursting. In order to determine the pausing index of RNA pol II, GRO-Seq data in mESCs was used (GEO ID: GSE48895). We obtained the fastq file from the bank (ENA accession number (fastq.gz): PRJNA 21123). As described previously, after removing the adapter sequence with the Cutadapt tool (version 2.4) (https://cutadapt.readthedocs.io/en/stable/index.html), reads were mapped to mm10 genome with Bowtie (Galaxy Version 1.1.2), and converted to bam file with SAM-to-BAM tool (Galaxy Version 2.1). This data were analyzed with the pausingIndex function of the groHMM tool (size = 500, up = 250, down = 250) (http://bioconductor.org/packages/release/bioc/html/groHMM.html) (version 1.10.0). Data of replication timing of mESCs was obtained from the following source (www.replicationdomain.org). Spearman’s rank correlation coefficient between either normalized TBi noise, burst size, or burst frequency and either promoter, or gene body localization degree (PRM) of various factors at the upper and lower 5% TCs of normalized TBi noise, burst size, and burst frequency was calculated. Next, the promoter-interacting distal enhancers were considered. Enhancers are believed to regulate gene expression by physical interaction with the promoter (*24*). Candidate distal *cis*-regulatory elements that interact with specific genes have been identified using capture Hi-C in mESCs (https://genomebiology.biomedcentral.com/articles/10.1186/s13059-015-0727-9). These data contain regions that interact with promoters and may include insulators and other elements in addition to enhancers. To identify the enhancers from regions that interact with the promoter of a particular gene, we manually screened for enhancers with relatively high RPM of H3K27ac ChIP-seq (RPM > 1.5)(Fig. S3L). Using this data, the RPM of other ChIP-seq data was calculated in the same manner as mentioned above in the candidate enhancers. Extreme outliers (with values 100 times the average) were excluded from the analysis. These enhancer data do not correspond to each TC, but rather correspond to each gene. Thus, the TBi noise, burst size, and burst frequency calculated using the gene-level count data were applied at this stage. Spearman’s rank correlation coefficient of normalized TBi noise, burst size and burst frequency with localization degree (PRM) of various factors in the upper and lower 5% enhancer of normalized TBi noise, burst size, and burst frequency of corresponding genes were calculated.

### OPLS-DA

Firstly, we classified promoter and gene body-associated features of high (either TBi noise, burst size, or burst frequency) transcripts into 10 clusters. Then, in order to identify the most contributing features for characterization of a cluster of high (either TBi noise, burst size, or burst frequency) transcripts against low (either TBi noise, burst size, or burst frequency) transcripts, we performed OPLS-DA modeling using ropls R package with 500 random permutations (version 1.8.0) (https://bioconductor.org/packages/release/bioc/html/ropls.html). One predictive component and one orthogonal component were used. To find the most influential variables for separation of high (either TBi noise, burst size, or burst frequency) groups against low (either TBi noise, burst size or burst frequency) groups, a S-plot with loadings of each variable on the x-axis and correlation of scores to modeled x-matrix (*p*(*corr*)[1]=*Corr*(*t1*,*X*), *t1* = scores in the predictive component) on the y-axis was constructed. Three each of the top and bottom variables with absolute value of loadings were selected.

### NGS and analysis of CRISPR library screening

After primer trimming with the Cutadapt software (https://cutadapt.readthedocs.io/en/stable/guide.html), read counts were generated and statistical analysis was performed using MAGeCK (v0.5.5) (https://sourceforge.net/p/mageck/wiki/Home/). DE scores were calculated from the gene-level significance returned by MAGeCK with the following formula: DE score = log_10_(gene-level depletion *P* value) – log_10_(gene-level enrichment *P* value). Genes with allelically normalized mean read count less than 10 from scRamDA-seq analysis were excluded from the downstream analysis. Then, genes were ranked by DE score. Subsequently, the top and bottom 100 genes were subjected to KEGG pathway analysis using an R package, clusterProfiler (v3.9.2) (https://github.com/GuangchuangYu/clusterProfiler).

### RNA degradation rate quantification

mRNA half-life can be determined using the following equation (33).

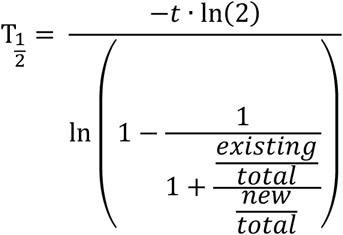

*t*, *existing*, *new*, *total* indicate the 4sU treatment time, amounts of existing, newly synthesized, and total RNA, respectively. Here, *t* is 1/3. *new*/*total* is 1-(*existing*/*total*).

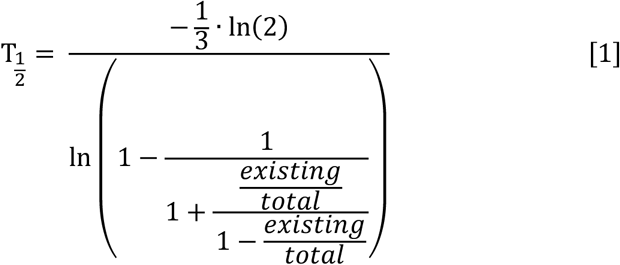

All samples contained spike-in RNA. Since they are unlabeled by 4sU and biotin, they are not trapped by streptavidin beads, except for nonspecific adsorption and technical loss. Therefore, by normalization with the amount of spike-in RNA in total and unbound (existing), the true ratio of total and unbound transcript can be obtained using the following equation.

Norm.Ratio(existing/total)=[unbound(target)/unbound(spike-in)]/[total(target)/total(spike-in)]=[unbound(target)/total(target)]/[unbound(spike-in)/total(spike-in)]

Unbound (target) / total (target) and unbound (spike-in) / total (spike-in) can be obtained by qPCR. Although most of the genes showed Norm.Ratio(existing/total) of more than 1, this is theoretically impossible (Fig. S6B). It is possible that reverse transcription efficiency is drastically decreased by biotinylation of RNA. Here, we assumed that the presence of biotinylated RNA during reverse transcription may trap reverse transcriptase, and that the efficiency of reverse transcription is further reduced globally. We assume that the global suppression effect of reverse transcriptase trapping is *I_g_* (global inhibitory effect). Moreover, the reverse transcription inhibitory effect of biotinylated RNA itself is defined as *I_s_*. Also, we defined *N*, *E*, *T*, and *R_eff_* as the amount of biotinylated (newly synthesized) RNA, the amount of existing unbiotinylated RNA, the amount of reverse transcriptase, and reverse transcription efficiency of reverse transcriptase, respectively. From these definitions, the cDNA amount derived from total and existing RNA can be determined by the following equations:

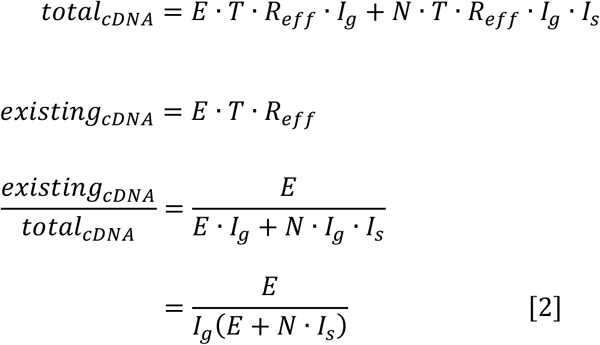

Next, a known value is introduced into the equation [1] to solve coefficients. The half-life of *Nanog* mRNA under Std conditions has been reported to be approximately 4.7 h (*18*). Therefore, the ideal ratio of existing / total *Nanog* mRNA amount is approximately 0.95203. In this case, the ideal relationship between newly synthesized and existing RNA is as follows.

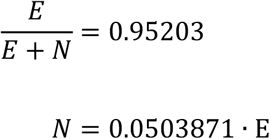

The mean ratio of existing/total *Nanog* cDNA revealed by qPCR was 3.436867. Therefore, the relationship between *I_s_* and *I_g_* is as follows from the equation [2].

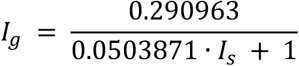

In order to determine the appropriate value of *I_s_*, several values were assigned to *I_s_*, and mRNA half-lives in the Std condition were compared with the previously reported mRNA half-lives (*21*) (Fig. S6C). We found that the scaling of mRNA half-lives in the Std condition and that of previously reported mRNA half-lives were quite similar when *I_s_* is 0.1 and *I_g_* is 0.289. Using equations [1] and [2], the half-lives of mRNA can be obtained based on the data using the value obtained from qPCR (Fig. S6D). No significant difference in mRNA half-life was observed between Std and PD-MK conditions for the genes examined.

## Acknowledgments

We would like to thank Tatsuo Miyamoto and Takashi Fukaya for helpful discussions, and Yuki Ochiai for technical assistance. We would like to also thank Prof. Joost Gribnau for providing the hybrid mouse ES cell line F1-21.6 (129Sv-Cast/EiJ, female). We thank Mr. Akihiro Matsushima and Mr. Manabu Ishii for their assistance with the infrastructure for the data analysis.

## Author contributions

Conceptualization, H.O.; Methodology, H.O., T.H., M.U., M.Y. and I.N.; Investigation, H.O., T.H., M.U., M.Y., I.N., A.H, Y.O., Y.S., K.N. and T.O.; Writing – Original Draft, H.O. and I.N.; Writing – Review & Editing, H.O., I.N., N.S., T.Y., H.K., Y.O. and Z.L.; Funding Acquisition, H.O. and I.N.; Supervision, H.O. and I.N.

## Data and materials availability

The datasets supporting the conclusions of this article are available in the GEO [GEO: GSE132593. This SuperSeries is composed of the SubSeries listed below: GSE132589, GSE132590, GSE132591, GSE132592.

## Supplementary Materials

**Figure S1.**
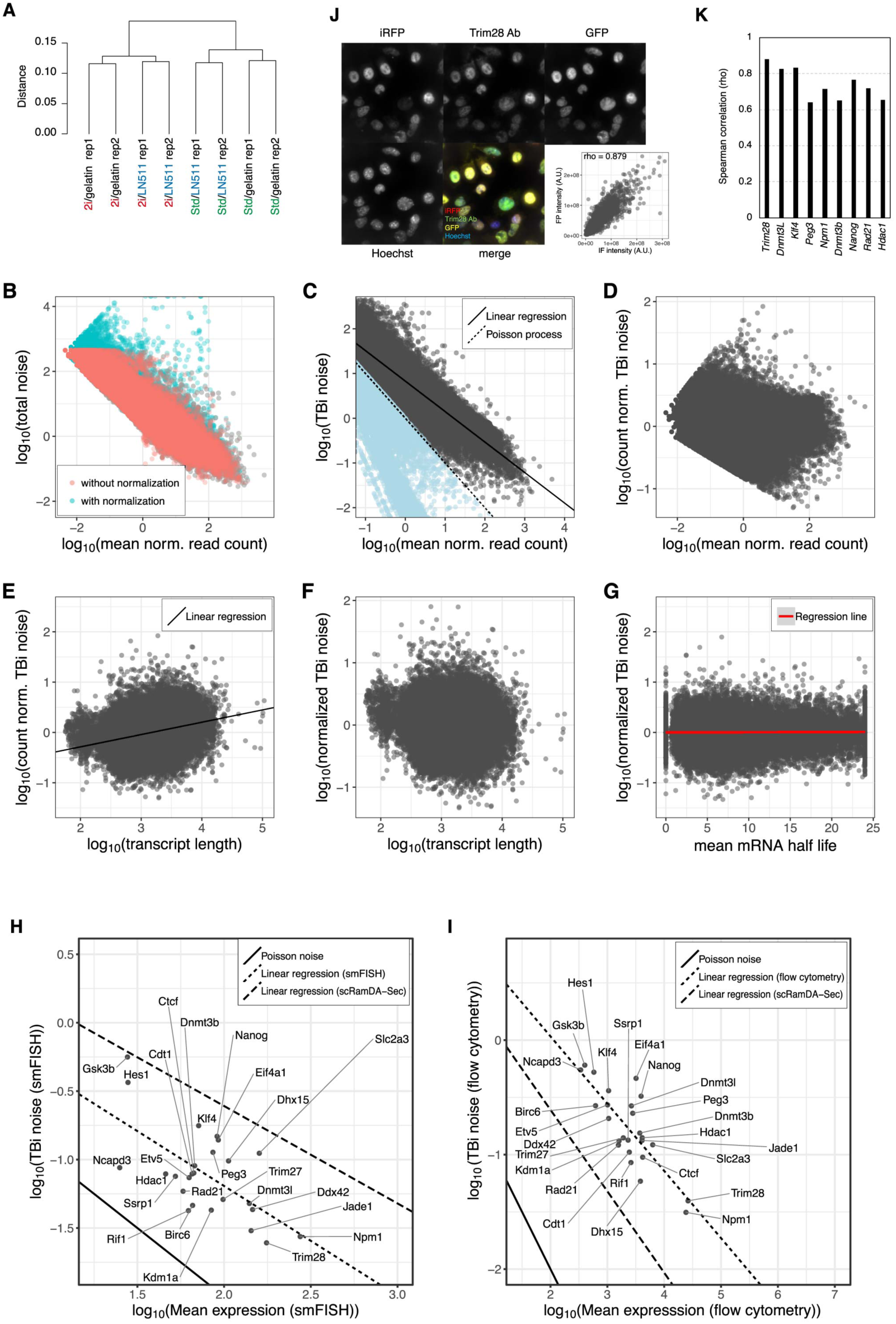
TBi noise determination using scRNA-seq data. **(A)** The difference between the transcriptomes of mESCs cultured on either Laminin-511 or gelatin coated dishes was relatively subtle. mESCs were conditioned on either gelatin or Laminin-511 (LN 511) coated dishes under either Std or 2i medium. RNA was extracted from these cells and analyzed by RNA-Seq. Two biological replicates were prepared. The result of clustering transcriptome data is shown. Biological replicates are similar to each other. The clusters are mainly divided into two clades depending on different medium conditions, suggesting that the influence of coating reagent difference (gelatin vs LN511) on the transcriptome is much smaller than that of the culture medium difference (Std vs 2i). **(B-G)** Processing of data obtained from scRNA-seq. **(B)** In order to calculate TBi noise, it was necessary to normalize so as to match the average expression level among alleles. Therefore, we first normalized the global expression level between alleles. Here, a scatter plot of the relationship between the mean read counts and total noise before and after a global normalization is shown. Even with global normalization, the shape of the distribution was not substantially changed. **(C)** Scatter plot of mean normalized read counts and TBi noise in data with expression levels that were normalized between alleles. Theoretically, TBi noise should not be less than Poisson noise (1 / mean read counts). Since data below the Poisson noise had a possibility of analytical defects, they were removed in subsequent analyses. TBi noise tends to decrease depending on the expression level. **(D)** Scatter plot of read count-normalized TBi noise and mean normalized read counts. **(E)** Scatter plot of count-normalized TBi noise and transcript length. **(F)** Scatter plot of count and transcript length-normalized TBi noise (referred as just normalized TBi noise) and transcript length. **(G)** Scatter plot of mRNA half-life and normalized TBi noise. There was no correlation between them (*r* = 0.0104). **(H-I)** Normalized TBi noise at mRNA level and protein level in KI cell line. To determine the normalized TBi noise at the mRNA and protein levels in the KI cell line, analysis was carried out with smFISH using allele specific probes **(H)** and flow cytometry **(I)** in these cell lines. Scatter plots of average expression level and TBi noise with regression lines are shown. The residuals from the regression line in the Y-axis direction were taken as normalized TBi noise. **(J-K)** Immunofluorescence of endogenous proteins in KI cell line. In some KI cell lines, the protein expressed from the target gene was fluorescently immunostained; the fluorescence image of immunofluorescence with GFP or iRFP derived from the knocked-in cassette was obtained by fluorescence microscopy. In addition, correlations of fluorescent intensity of individual cells were examined. Since the KI cassette contains 2A peptide, nuclear localization signal (NLS)-GFP (or iRFP), and PEST sequence, which induces rapid protein degradation, these endogenous proteins and knocked-in fluorescent proteins become different protein molecules after translation. **(J)** Fluorescent images of fluorescent proteins and immunostained endogenous TRIM28 proteins in the *Trim28* KI cell line. The images show maximum intensity projections of stacks. The lower right panel shows a scatter plot of the fluorescence intensities of the fluorescent proteins and the fluorescent immunostaining. **(K)** A bar graph of the Spearman’s rank correlation coefficients between the fluorescence intensities of the fluorescent protein and the fluorescent immunostaining of the endogenous protein. All proteins analyzed here showed substantial correlation, suggesting that the fluorescence intensity of the fluorescent protein is indicative of endogenous protein abundance.

**Figure S2.**
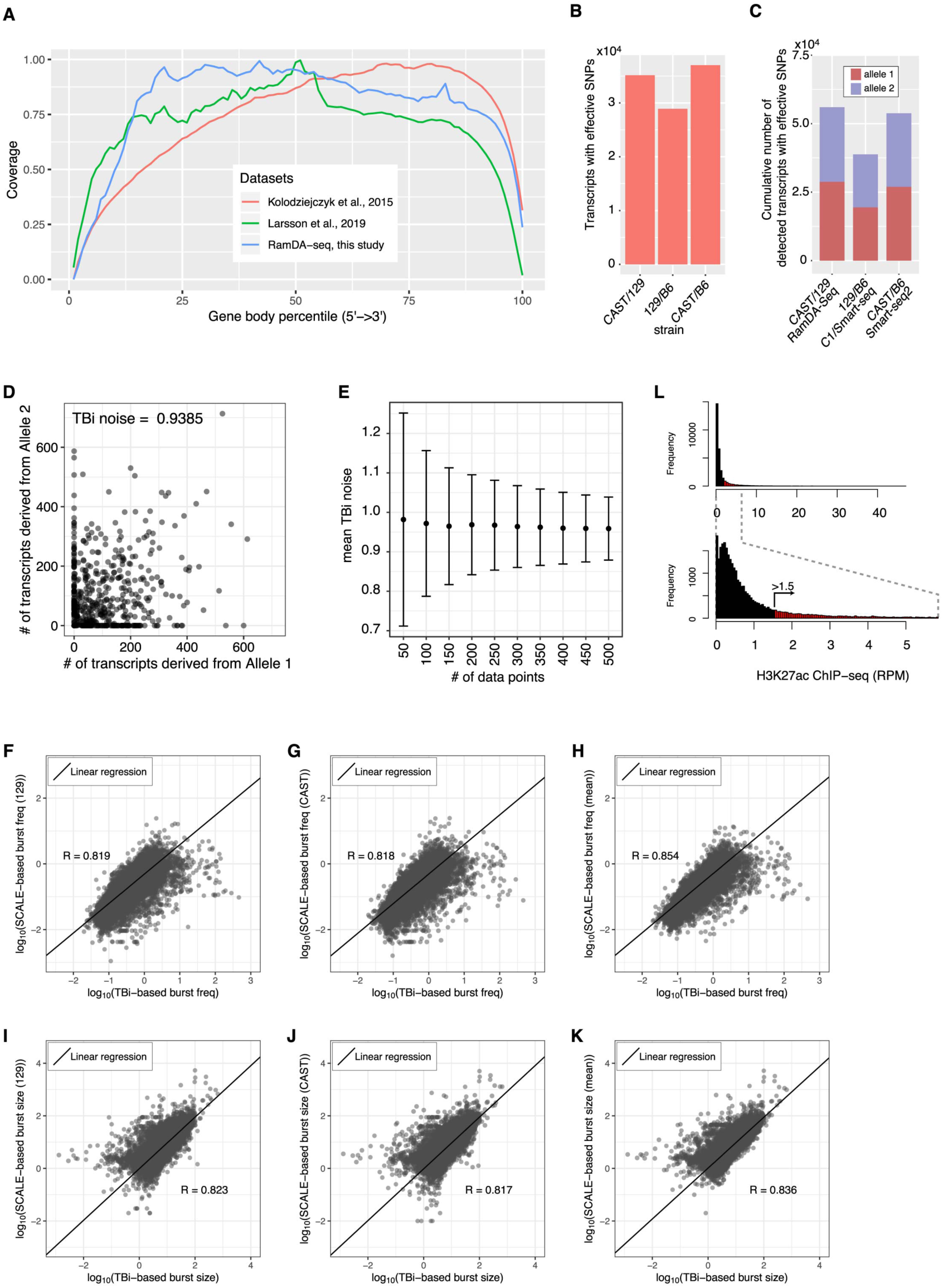
Comparison of the classification efficiency of each allele using SNPs with results of single-cell (sc) RNA-seq (RamDA-seq) with that of other studies. **(A)** Mean read coverage over transcripts of three datasets. “RamDA-seq, this study”: CAST/129 mESC scRamDA-seq; “Kolodziejczyk et al., 2015”: 129/B6 mESC C1/Smart-seq performed in (*36*); “Larsson et al., 2019”: CAST/B6 mESC Smart-seq2 performed in (*8*). **(B)** The theoretical numbers of transcripts with effective SNPs in three different hybrid mESC genomes. **(C)** The cumulative number of detected transcripts with effective SNPs in three different experiments. CAST/129 RamDA-seq: CAST/129 mESC scRamDA-seq in this study; 129/B6 C1/Smart-seq: 129/B6 mESC C1/Smart-seq performed in (*36*); CAST/B6 Smart-seq2: CAST/B6 mESC Smart-seq2 performed in (*8*). The classification efficiency of each allele using the SNPs of this study was comparable to that of the other studies. **(D, E)** The number of samples affects the confidence of calculated TBi noise. **(D)** Each of the 1000 simulated datasets with 500 data points of the set of allele expression was generated. Mean expression of each allele and TBi noise in these data sets were 113.22±5.74 (SD) and 0.958±0.0799 (SD), respectively. One representative plot is shown. **(E)** Differences in standard deviation between datasets with different numbers of data points. We calculated the mean and standard deviation of TBi noise in datasets with different numbers of data points. Error bars indicate standard deviation. It is evident that the larger the number of data points, the smaller the data variation. We used 447 of 129/CAST mESCs in the G1 phase for the analysis of TBi noise in this study, while, Kolodziejczyk et al., and Larsson et al. used 250 of 129/B6 mESCs, and 188 of B6/CAST mESCs that were in different phases of the cell cycle and were cultured in Std medium or unknown condition in their scRNA-seq, respectively (*8, 36*). Thus, our data are expected to be an important resource for a deeper understanding of transcriptional bursting and TBi noise. **(F-K)** Comparison of burst frequencies **(F-H)** and sizes **(I-K)** inferred using either TBi noise levels or SCALE software (see Material and Methods). By using SCALE, the burst size and frequency parameters of individual alleles can be determined. **(F, I)** Scatter plots of TBi noise-based vs. SCALE-based (129 allele) parameters with regression lines. **(G, J)** Scatter plots of TBi noise-based vs. SCALE-based (CAST allele) parameters with regression lines. **(H, K)** Scatter plots of TBi noise-based vs. SCALE-based (mean of CAST and 129 allele values) parameters with regression lines. **(L)** Histogram of reads per million reads (RPM) of H3K27ac ChIP-seq of promoter-associated regions in mESCs (see Material and Methods). Enhancers were manually defined as H3K27ac ChIP-seq RPM greater than 1.5.

**Figure S3.**
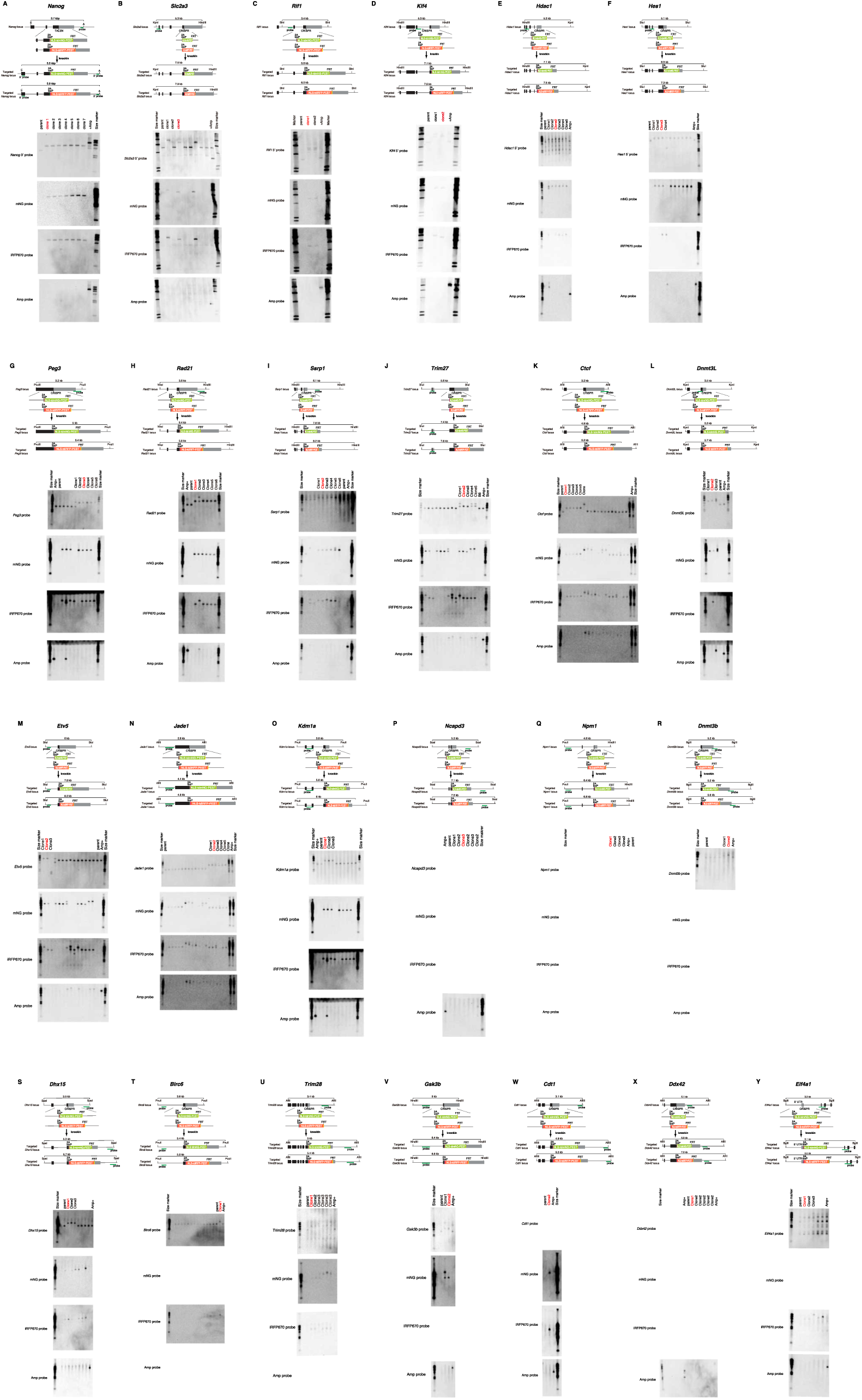
Establishment of knock-in cell lines. The upper part of the panel shows structures of the target gene near the knock-in site, the targeting vector, and the knocked-in genes. The results of Southern blotting are shown at the bottom of the panel. Parent means the parental strain (C57BL6J, Bruce4), +Amp means a cell line in which the ampicillin resistance gene, which is contained in the backbone of the targeting vector, has been introduced into the genome. Cell lines shown in red letters were used in the downstream experiments. See Table S6 for details of knock-in cell lines used in this study.

**Figure S4.**
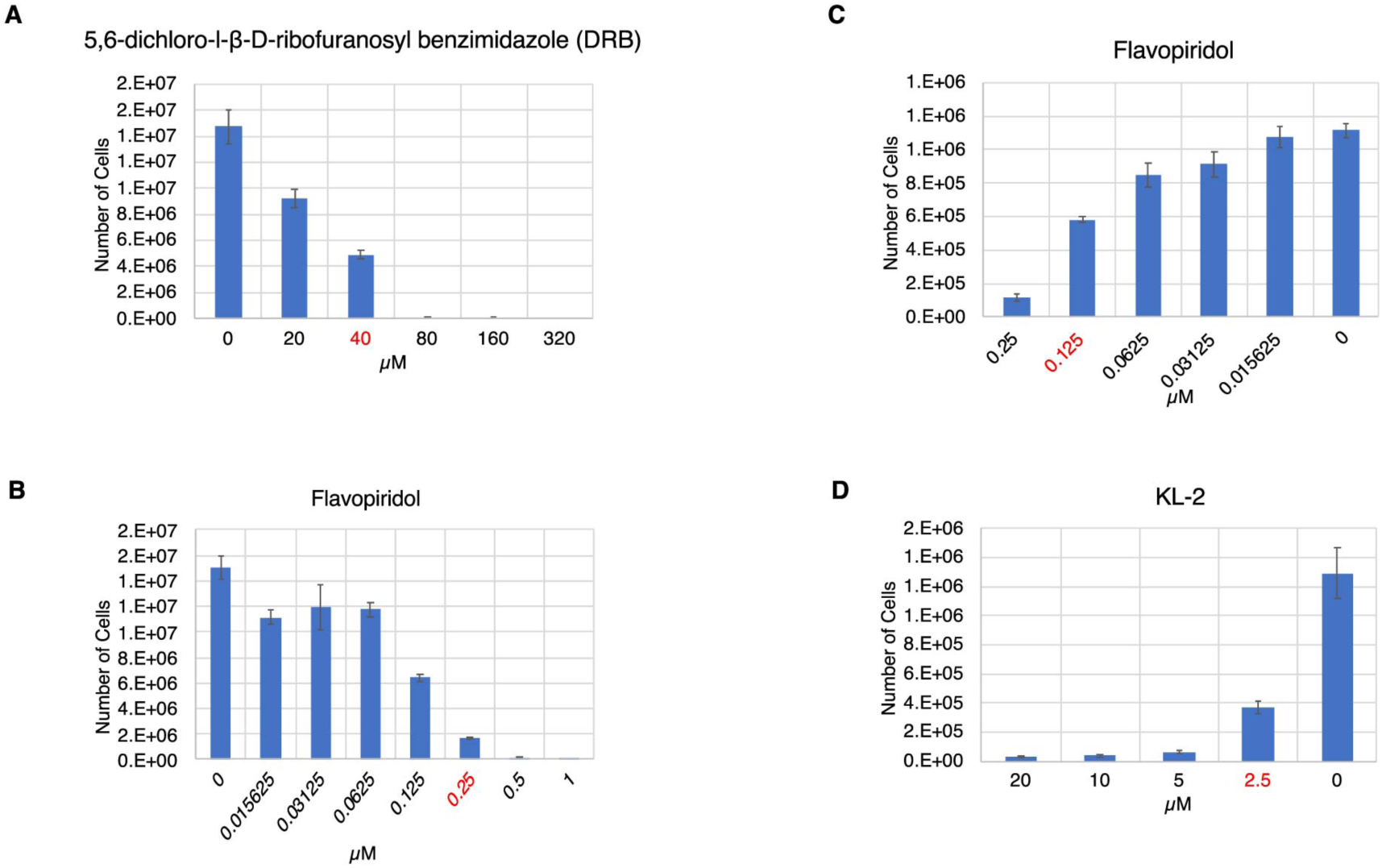
Effects of transcription elongation inhibitor on cell growth. **(A, B)** 1 × 10^5^ C57BL6J WT mESCs conditioned to 2i medium were cultured for 2 days in the presence of 5,6-dichloro-l-β-D-ribofuranosyl benzimidazole (DRB) **(A)** or Flavopiridol **(B)**, and the number of cells were counted. We decided to use the concentration (highlighted in red letters), at which the growth rate drops sufficiently, and cells were not extinct, in the following experiment. **(C, D)** Effects of transcription elongation inhibitor on cell growth in mESCs conditioned PD-MK medium. 1 × 10^5^ WT mESCs conditioned to PD-MK medium were cultured for 2 days in the presence of Flavopiridol **(C)** or SEC inhibitor KL-2 **(D)**, and the number of cells were counted. We decided to use the concentration (highlighted in red letters), at which the growth rate drops sufficiently and cells were not extinct, in the following experiment.

**Figure S5.**
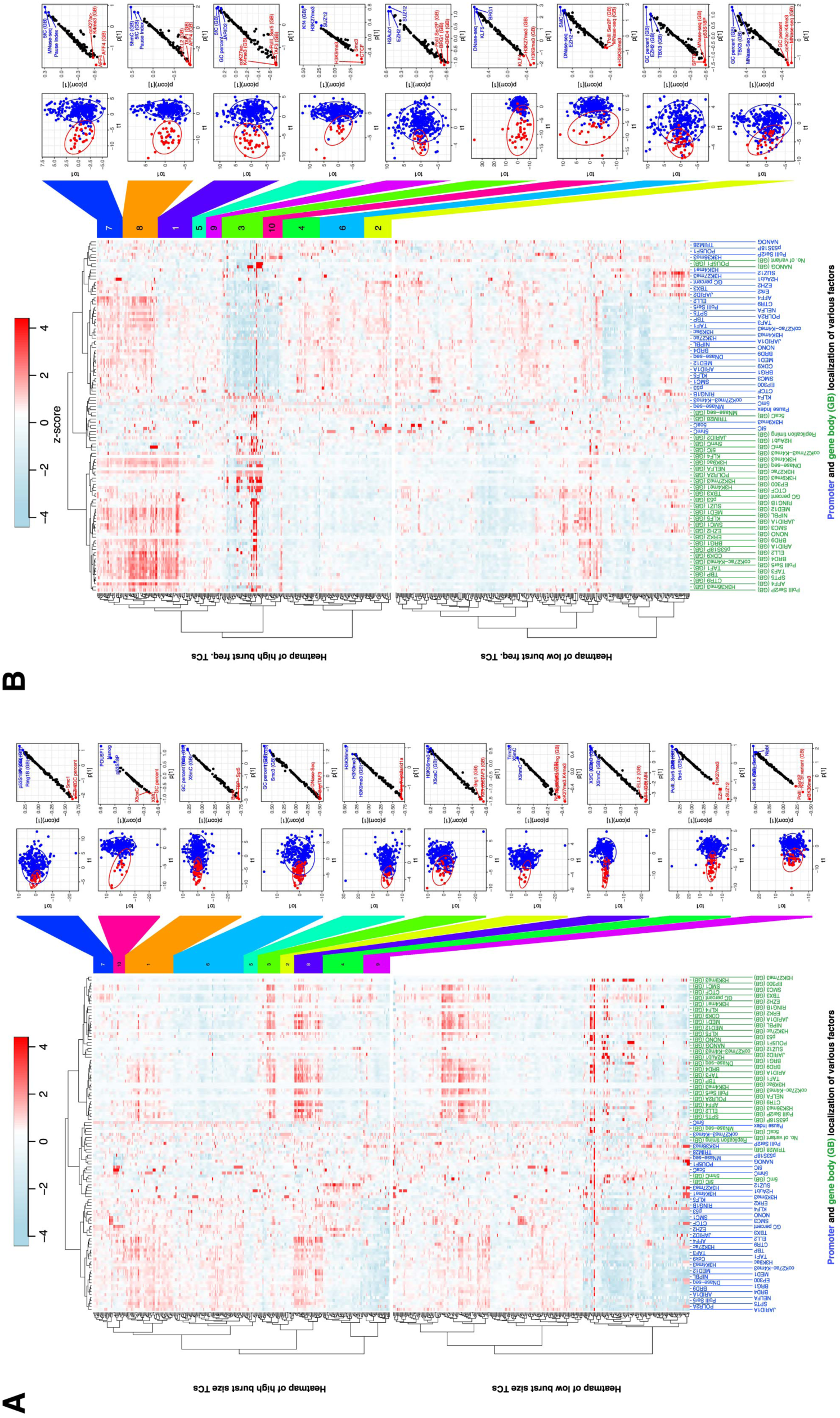
Burst size and frequency are determined by combinations of promoter and gene body binding factors. **(A)** Burst size is determined by combinations of promoter and gene body associating factors. First, the target 5992 transcripts (TCs) were ranked with the burst size, of which the upper or lower 5% (300 TCs each) was taken as high and low burst size TCs, respectively. The left side of the panel shows a heat map of promoter and gene body (GB) association of various factors in high and low burst size TCs. The high burst size TCs were classified into 10 clusters, and each cluster of high burst size TCs and whole low burst size TCs were subjected to OPLS-DA modeling. The right side of the panel represents score plots of OPLS-DA and S-plots constructed by presenting covariance (*p*) against correlation [*p*(*corr*)]. **(B)** Burst frequency is determined by combinations of promoter and gene body associating factors. First, the target 5992 transcripts (TCs) were ranked with the burst frequency, of which the upper or lower 5% (300 TCs each) was taken as high and low burst frequency TCs, respectively. The left side of the panel shows a heat map of promoter and gene body (GB) association of various factors in high and low burst frequency TCs. The high burst frequency TCs were classified into 10 clusters, and each cluster of high burst frequency TCs and whole low burst frequency TCs were subjected to OPLS-DA modeling. The right side of the panel represents score plots of OPLS-DA and S-plots constructed by presenting covariance (*p*) against correlation [*p*(*corr*)].

**Figure S6.**
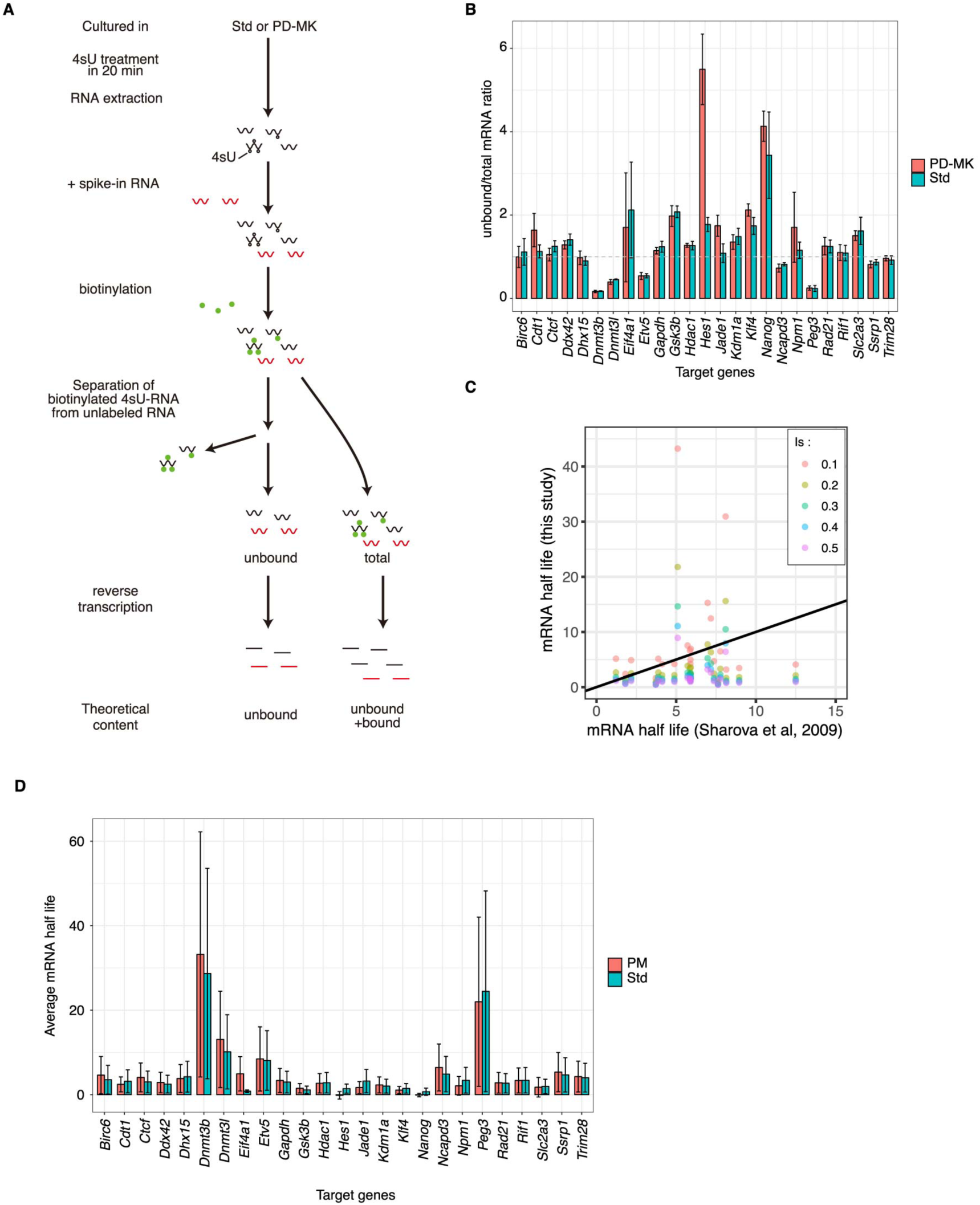
RNA degradation rate between PD-MK and Std conditions did not show significant difference. **(A)** Schematic representation of the experiment for examining RNA degradation rate. WT mESCs were transiently treated with 4-thiouridine (4sU). By this transient treatment, 4sUs were incorporated into newly synthesized RNA. RNA was then extracted from the cells, and a known amount of spike-in RNA that does not contain 4sU was added. Thereafter, the RNA mix was biotinylated. Then, from a part of this biotinylated RNA mix, biotinylated RNA was removed using streptavidin beads, and unbiotinylated RNAs that were transcribed before the addition of 4sU were recovered. RNA samples that were not treated with streptavidin beads contained both existing RNA and newly synthesized RNA. Therefore, we refer to these RNA samples as total RNAs. These were reverse transcribed and analyzed by qPCR. **(B)** A bar graph of the ratio of unbound and total RNA. For many samples, the ratio has exceeded 1, which was theoretically impossible. It is considered that the reverse transcription efficiency of 4sU-introduced RNA could be extremely low (see Material and Methods). **(C)** We assumed that the presence of biotinylated RNA during reverse transcription may trap reverse transcriptase, and that the efficiency of reverse transcription is further reduced globally. We assume that the global suppression effect of reverse transcriptase trapping is *I_g_* (global inhibitory effect). Moreover, the reverse transcription inhibitory effect of biotinylated RNA itself was defined as *I_s_* (see Material and Methods). In order to determine the appropriate value of *I_s_*, several values were assigned to *I_s_*, and mRNA half-lives in the Std condition were compared with the previously reported mRNA half-lives (*21*). We found that the scaling of mRNA half-lives in the Std condition and that of previously reported mRNA half-lives were getting closer when *I_s_* was 0.1. **(D)** Average mRNA half-life. The half-life of mRNA was calculated using the ratio obtained in **(B)** and the correction formula (see Material and Methods). In all genes analyzed, there was no significant difference between PD-MK and Std conditions.

---

**Table S1. Allelically normalized read count data of individual transcripts of 129 alleles.** Data with TBi noise below Poisson noise or transcripts showing interallelic extreme expression level differences are removed (see Material and Methods).

**Table S2. Allelically normalized read count data of individual transcripts of CAST alleles**. Data with TBi noise below Poisson noise or transcripts showing interallelic extreme expression level differences are removed (see Material and Methods).

**Table S3. smFISH count data of GFP and iRFP knocked-in allele in knock-in cell line.**

**Table S4. Plasmids used for knock-in cell line establishment, sequences of oligos used in this study, and list of data sources used in this study.**

